# A Chemoproteomic Atlas of the Human Purine Interactome for Regioselective Ligand Discovery

**DOI:** 10.1101/2025.10.21.683656

**Authors:** Zhihong Li, Hsiao-Kuei Tsai, Adam H. Libby, Michael W. Founds, Olivia L. Murtagh, Madeleine L. Ware, David M. Leace, Wesley J. Wolfe, Phillip W. Gingrich, Bissan Al-Lazikani, Chin-Yuan Chang, Ku-Lung Hsu

**Author notes:** Author to whom correspondence should be addressed: (K.-L.H.) Department of Chemistry, University of Texas at Austin, 100 E 24th St, NHB 6.406, A5300, Austin, Texas 78712.

## Abstract

Purines are essential bioactive molecules that interact with a large fraction of the human proteome. Despite their importance, the scope of actionable purine-binding pockets for ligand discovery remains limited. Here, we developed a quantitative chemoproteomics platform using sulfonyl-purine (SuPUR) chemistry to produce a massive and functional map of the human purine interactome. The SuPUR platform captured 31,000+ targetable tyrosine and lysine sites, representing the most comprehensive beyond cysteine chemoproteomics database for enabling protein ligand discovery. SuPUR ligands that bind through a regioselective fashion serve as enabling starting points for developing potent (nanomolar) and proteome-wide-selective modulators of enzymatic and protein-protein interaction function. Phenotypic screening identified a site-specific (Y237) and regioselective SuPUR ligand of ACAT2 to reveal an unexpected metabolic dependency in cancer cells. A crystal structure of SuPUR ligand-bound ACAT2 revealed the purine group binds deep in the CoA pocket forming key interactions with catalytic residues via a water bridge to guide future structure-based ligand design.

---

Purine is a ubiquitous metabolite found in nature and serves as a building block for producing key bioactive compounds involved in diverse cellular functions^1–4^. Substituted purines are components of energy cofactors (adenosine triphosphate (ATP) and guanosine triphosphate (GTP)), coenzymes in oxidation-reduction reactions, secondary messengers (cyclic AMP, cyclic GMP), neurotransmitters (adenine), enzyme cofactors (nicotinamide adenine dinucleotide and flavin adenine dinucleotide), inflammatory signals (urate), and nucleotides (**Fig. 1a**). Purine levels in mammalian cells are maintained through the coordinated regulation of biosynthetic and salvage pathways.^5, 6^ The human genome is predicted to encode ∼3,000 proteins that bind purines although the full repertoire of purine-binding proteins likely extends beyond what can be predicted by sequence homology alone.^7–9^

**Fig. 1.**
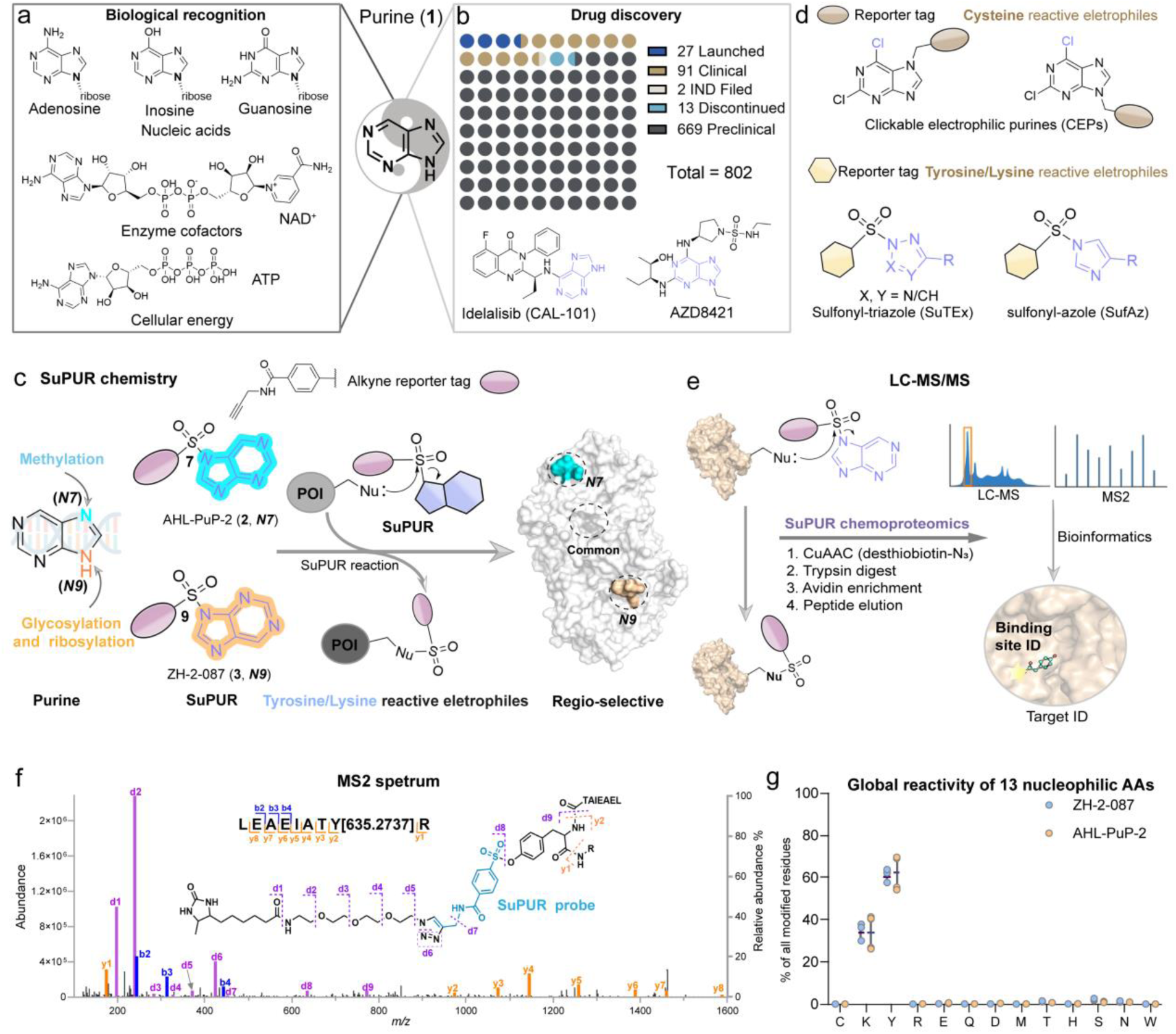
Developing SuPUR chemistry to target the human purine interactome. **a,** Purines include adenine and guanine, which are essential blocks for the formation of DNA and RNA. Additionally, purines are key components of various important biomolecules, such as ATP, GTP, cyclic AMP, NAD(P)+, and coenzymes. **b,** Purine (**1**) serves as a privileged scaffold in drug discovery, with increasing clinical drugs incorporating the purine scaffold. **c**, Development of sulfonyl-purine (SuPUR) probes AHL-PuP-2 (**2**) and ZH-2-087 (**3**) that are proposed to mediate recognition via the purine leaving group. The purine core enables the installation of a sulfonyl electrophile at the N7 or N9 position for evaluating regioselective covalent binding to nucleophilic residues on proteins of interest (POI). **d,** Clickable electrophilic purine (CEP) probes were designed for covalent targeting of cysteine residue, and sulfonyl-containing chemical handles were employed to target tyrosine and lysine residues selectively.^27, 31^, _32_ **e,** SuPUR chemoproteomics to investigate the targetable human purine interactome. **f,** Representative MS2 spectrum annotation of AHL-PuP-2 modified site (Y380) found in KRT18 (UniProt: P05783). (n = 3). **g,** The global reactivity of 13 nucleophilic amino acids (AAs) with AHL-PuP-2 (N7-isomer) and ZH-2-087 (N9-isomer), respectively. Data shown are mean ± SD (n = 3). More examples are provided in Extended Data Fig. 1.

Efforts to study the purine interactome include development of chemical probes that incorporate modified purines as recognition elements to facilitate protein binding. Specifically, in chemoproteomics, the purine moiety is a key component of ATP acyl phosphate probes for activity-based protein profiling (ABPP) of kinases and other ATP-binding proteins in proteomes.^10, 11^ The adenine portion of these ATP substrate mimetic probes is important for mediating molecular interactions in the kinase active site and align the acyl phosphate electrophile for covalent capture of conserved catalytic lysines. Adenine is key for recognition and modification of this group to other purine analogs changes the resulting interactome (e.g., ATP- vs GTP-based probes to bind ATPases and GTPases, respectively).^12^ Additional purine-based targeting strategies include chemoproteomic probes incorporating a coenzyme A (CoA) recognition element to investigate lysine acetyltransferases or photoreactive *S*-adenosyl homocysteine (SAM) analogs for ABPP analysis of methyltransferases.^13–18^

Purine (**1**) is found in drugs and considered a privileged scaffold due to its ability to bind a broad range of protein targets, with examples of drugs targeting PI3K (idelalisib) and the recently disclosed CDK2 inhibitor AZD-8421.^19, 20^ Importantly, an increasing number of preclinical and clinical drugs incorporating the purine core are rapidly emerging (**Fig. 1b**). While specific purine-based drugs are reported, purine-binding pockets in general are challenging to target selectively because of the intrinsic functional redundancy. For example, ATP- and NADP-binding pockets are highly conserved between functionally distinct protein classes.^21, 22^

The nature and position of chemical modifications on the purine core can mediate biological specificity. The oxygenation and amination patterns at the C2 and C6 positions of purine are critical for proper base pairing of DNA.^23, 24^ The purine N7 and N9 positions can be modified regioselectively to change function and fate of the resulting biomolecules. Glycosylation and ribosylation of N9 is enzymatically preferred for nucleoside biosynthesis while methylation at N7, and on guanosine specifically, can alter mRNA stability and translation (**Fig. 1c**).^25, 26^ Purine recognition sites, therefore, are promising targets for ligand and drug development if technologies to systematically evaluate ligandability were readily available to guide discovery efforts.

Recently, clickable electrophilic purines (CEPs) were deployed to investigate RNA-binding protein (RBP) activity in cells. CEPs covalently modify RBPs and other purine-binding proteins via nucleophilic aromatic substitution reaction between cysteine and the electrophilic C6 position (**Fig. 1d**).^27^ To enable chemoproteomic detection, the N7 or N9 positions were modified with an alkyne group to facilitate click chemistry conjugation of reporter tags for detection of CEP-modified proteins by gel- or liquid chromatography-mass spectrometry (LC-MS/MS)-based readouts, respectively. These findings support the purine core itself as a suitable recognition element and provides rationale for modifying the electrophile to target residues beyond cysteine and expand the scope of purine-binding pockets captured by chemoproteomics (**Fig. 1c**, **d**).

Here, we deployed a quantitative and multiplexed chemoproteomics platform using sulfonyl-purine (SuPUR) chemistry to produce a massive and functional map of the human purine interactome. Deep SuPUR chemoproteomic profiling of >30,000 distinct tyrosine and lysine sites – representing the most comprehensive beyond cysteine chemoproteomic database – revealed the human proteome is replete with purine binding pockets as candidate targets for ligandability assessment. The SuPUR scaffold enabled rapid development of potent (nanomolar) and proteome-wide-selective ligands that regioselectively engage NAD+ and CoA binding pockets to disrupt enzymatic and protein-protein interaction functions. Phenotypic screening revealed a targetable dependency in cancer cells through site-specific and regioselective SuPUR binding to ACAT2 Y237, a key enzyme in cholesterol biosynthesis. A co-crystal structure revealed that the ACAT2 ligand binds deep in the CoA pocket, forming key interaction between purine and catalytic residues via a water bridge to help support the observed regioselectivity.

## Results

### Development of SuPUR chemoproteomics

SuPUR probes were synthesized by sulfonylation of the purine N7 or N9 with a minimal alkynyl reporter design previously demonstrated to be compatible with LC-MS/MS analyses.^28^ We speculated the purine exhibits sufficient leaving group ability to activate SuPUR modification of proteins in a manner analogous to reported sulfonyl-azole chemistry (**Fig. 1d**).^29, 30^ Strategic installment of the sulfonyl electrophile at the N7 vs N9 position affords opportunities for assessing regioselective recognition at individual protein sites (**Fig. 1c**).

To assess protein labeling activity, HEK293T cell proteomes were treated with the SuPUR probes AHL-PuP-2 (N7 isomer, **2**) or ZH-2-087 (N9 isomer, **3**) followed by click chemistry (CuAAC^33^) conjugation of rhodamine-azide and SDS-PAGE detection of fluorescent protein bands. These gel-based chemoproteomic studies demonstrated comparable concentration- and time-dependent labeling activity for the SuPUR probes (Extended Data Fig. 1a-d). Importantly, pretreatment with free purine resulted in a substantial and concentration-dependent loss of AHL-PuP-2 probe labeling activity (Extended Data Fig. 1e-g). Competition with benzimidazole (**4**), which is similar to purine but lacks key nitrogen atoms required for purine recognition, demonstrated a lack of inhibitory activity in the probe competition study. These findings support purine recognition as an important factor for the observed SuPUR probe labeling in proteomes.

Next, we conducted label-free, LC-MS/MS chemoproteomics to identify nucleophilic residues on proteins preferentially modified by SuPUR chemistry (**Fig. 1e**). Careful inspection of MS2 spectra from SuPUR probe-modified peptides identified y- and b-fragment ions that confidently localized probe modifications on residues modified by the SuPUR reaction (SuPUR adduct mass of 635.2737 Da, **Fig. 1f** and **Methods**). Across the 13 nucleophilic amino acids evaluated on detectable peptides, we determined AHL-PuP-2 and ZH-2-087 preferentially modified tyrosine (Y, ∼60%) and lysine (K, ∼40%; **Fig. 1g**). Compared with SuTEx,^34^ SuPUR chemistry appeared to show higher lysine binding preference (40% vs 25% K modifications, respectively; **Fig. 2a**). Binding site coverage between AHL-PuP-2 and ZH-2-087 was comparable in SuPUR probe-modified HEK293T proteomes (∼14,000-15,000 probe-modified sites, **Fig. 2a**).

**Fig. 2.**
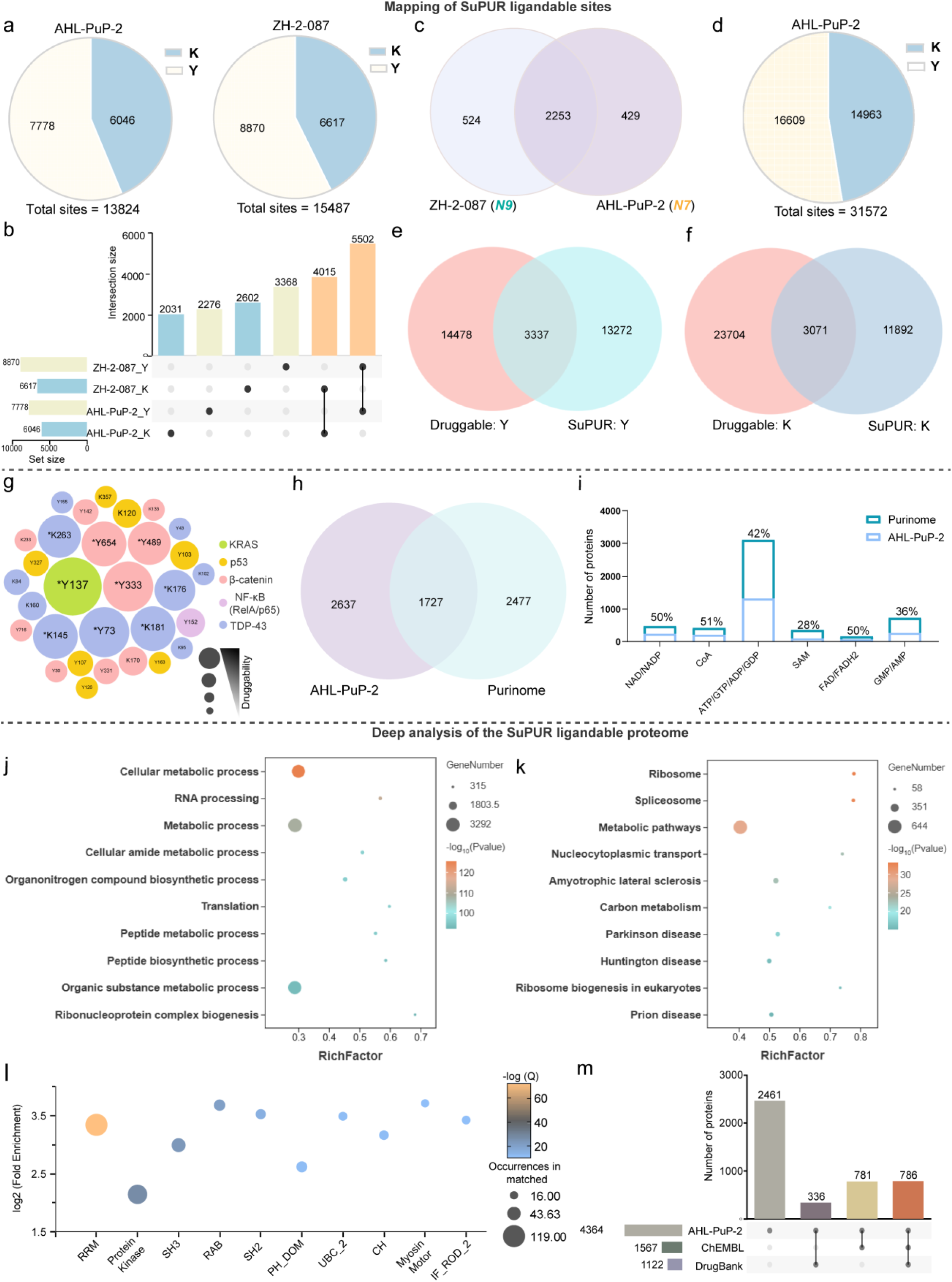
SuPUR chemoproteomics establishes the targetable human purine interactome. **a,** AHL-PuP-2 (N7-isomer) and ZH-2-087 (N9-isomer) modified tyrosine (Y) and lysine (K) sites were identified in HEK293T proteome (100 µM probe, 1 h, rt). The sites represent the combined results from n = 3 independent biological replicates. **b** and **c,** Comparison of AHL-PuP-2 and ZH-2-087 modified sites (**b**) and proteins (**c**) from analyses in **a**. **d,** Aggregate of AHL-PuP-2 (100 µM, 1 h, rt) modified tyrosine (Y) and lysine (K) sites detected in proteomes from human cells (HEK293T, A549, MDA-MB-231, Colo-205, SiHa, and Jurkat cells). The sites represent the combined results from n = 3 independent biological replicates. **e** and **f**, Comparison of predicted druggable tyrosine (Y; **e**) or lysine (K; **f**) with AHL-PuP-2 modified tyrosine or lysine sites, respectively. **g,** Representative sites targeted by SuPUR chemoproteomics on intractable proteins of interest. Asterisk denotes sites located in a predicted druggable pocket. **h**, Comparison of proteins targeted by AHL-PuP-2 with the human purinome. See Supporting Methods for identification of purine-binding proteins from available public datasets. **i,** Analysis of human purinome subclasses that are targeted by SuPUR chemoproteomics (AHL-PuP-2). **j,** Gene ontology (GO) enrichment analysis of AHL-PuP-2 modified proteins for biological process; the top 10 enriched GO terms are shown. **k,** KEGG pathway enrichment analysis of AHL-PuP-2 identified proteins; the top 10 enriched KEGG terms are shown. **l,** Domain enrichment analysis of SuPUR-modified sites. See Supporting Methods for details of domain enrichment analyses. **m,** Comparative analysis of protein targets identified by AHL-PuP-2 with DrugBank and ChEMBL databases. More examples are provided in Extended Data Fig. 2.

Comparison of probe-modified sites (Y and K) labeled by each respective probe revealed overlapping (∼64%) but also, interestingly, distinct sites for the N7- (AHL-PuP-2, 31%) versus N9-isomer probes (ZH-2-087, 38%) that also translated into distinct protein targets (**Fig. 2b**, **c**). Proteins labeled by N7-compared with N9-SuPUR probes were enriched for different biological functions as measured by Gene Ontology analyses (GO analyses, Extended Data Fig. 2a, b).^35, 36^ Furthermore, we detected differences in the functional domains enriched by the N7- vs N9-SuPUR probes (Extended Data Fig. 2c, d). These data highlight the capability of SuPUR probes for regioselective and proteome-wide detection of targetable protein sites.

We expanded chemoproteomic profiling to 5 additional human cell proteomes for determining the breadth of proteomic coverage attainable with the SuPUR platform. Using AHL-PuP-2, we detected on average ∼13,000 modified sites from each respective cell line that in aggregate translates to 31,000+ total detected sites (∼50/50 distribution between Y and K sites; referred herein as the SuPUR interactome) from ∼4,300 proteins (**Fig. 2d** and Supplementary **Table 1**). To the best of our knowledge, this chemoproteomic dataset represents the most comprehensive database of covalently targetable tyrosine and lysine residues in the human proteome. We discovered that ∼20% of SuPUR interactome sites reside in a predicted drug pocket, providing insights to which detectable sites are likely candidates for further ligand development (**Fig. 2e**, **f**). We identified several emerging (b-catenin Y333, Y489, Y654; NF-kB Y152; TDP-43 K145, K176) and known clinical targets (KRAS Y137; p53 K120) that possessed at least one SuPUR probe-modified site in a predicted druggable pocket (**Fig. 2g** and Extended Data Fig. 2e-i).^37, 38^

Importantly, the SuPUR interactome captures a substantial fraction (41%) of the human purine interactome with coverage across multiple subclasses of purine-binding proteins (30-50%; **Fig. 2h**, **i**). Finally, we conducted GO and KEGG pathway enrichment analyses of the SuPUR interactome proteins to investigate their associated biological processes.^35, 36, 39^ The top enrichments included cellular metabolism and RNA processing, which is supported by the role of purine as an endogenous metabolite (e.g., ATP, GTP, NAD^+^, adenosine) and a building block for RNA (**Fig. 2j**, **k**; Supplementary **Table 2**).

### Deep analysis of the SuPUR ligandable proteome

We performed domain enrichment analysis on SuPUR interactome sites to gain a deeper understanding of protein domains targeted.^40^ As expected, we observed enrichment for RNA-binding regions (RRM domain), which matches the known role of purine as a core constituent of nitrogenous bases found in RNA (**Fig. 2l**). Beyond RNA, domains involved in binding ATP (protein kinase domain) and GTP (RAB domain), which are cofactors that contain a purine component were also enriched in the SuPUR interactome dataset. To determine whether SuPUR interactome proteins have known drugs or ligands, we performed a comparative analysis of AHL-PuP-2-modified proteins against DrugBank and ChEMBL databases.^41, 42^ The analysis revealed that 56% of SuPUR interactome proteins lack known bioactive ligands or approved drugs, supporting SuPUR chemoproteomics as an enabling technology for developing ligands against historically intractable proteins (**Fig. 2m**).

To assess ligandable regions on SuPUR interactome proteins, we synthesized ZH-1-049-2 (**5**) and ZH-2-055 (**6**) and confirmed by X-ray crystallography the N7- and N9-regioisomeric state, respectively, for evaluation in a competitive tandem mass tag (TMT)-ABPP assay (**Fig. 3a**, **b**; Supplementary **Table 3**).^32^ To assess purine recognition, the matching single nitrogen deletion (N-delete) control compounds ZH-2-036 (N7, **7**) and ZH-5-019 (N9, **8**) were synthesized and included for direct comparison (**Fig. 3c**, **d**). The regiochemically defined structures for this SuPUR fragment ligand set were further confirmed by carbon (^13^C) NMR analysis, which revealed detectable differences in the C4/C5 and C5 signals of the SuPUR and N-delete regioisomer pairs, respectively (**Fig. 3b**, **d**).

**Fig. 3.**
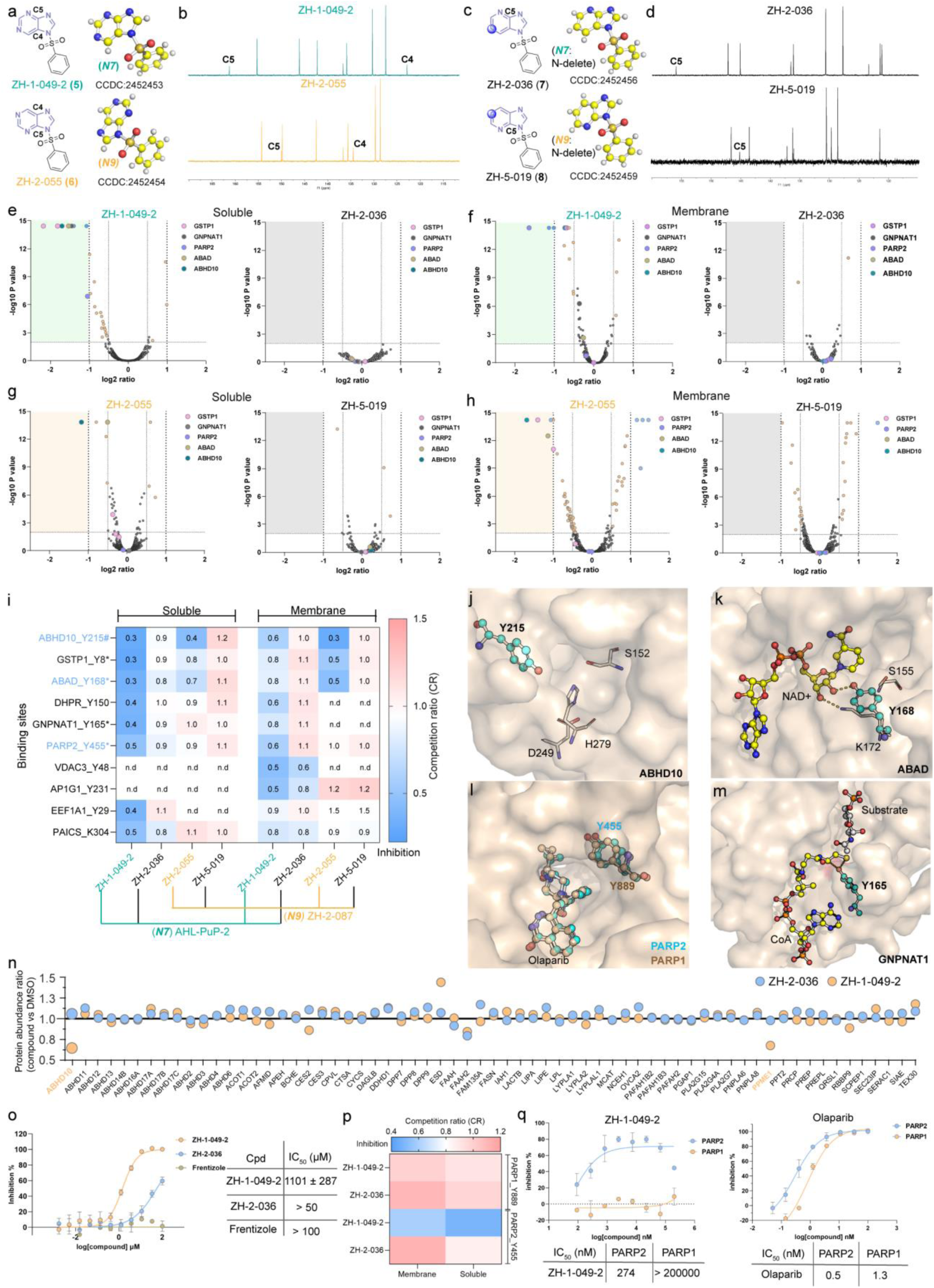
Lead SuPUR ligands demonstrate site-specific and regio-selective protein modulation. **a,** Development of N7-isomer SuPUR ligand ZH-2-049-2 (**5**), and the matching N9-isomer ZH-2-055 (**6**). Crystal structures confirming regioisomer pair structures are shown. **b,** (^13^C) NMR data for ZH-1-049-2 and ZH-2-055 showed significant chemical shifts in C5 and C4 signals between the N9- and N7-substituted products. **c,** Development of matching control molecules ZH-2-036 (**7**) and ZH-5-019 (**8**) by deleting a nitrogen on the purine core, along with their corresponding crystal structures. **d,** (^13^C) NMR data for ZH-2-036 and ZH-5-019 showed significant chemical shifts in the C5 signal between the N9- and N7-substituted products. **e** and **f,** Competitive SuPUR TMT-ABPP analysis of ZH-1-049-2 and ZH-2-036 (25 µM, 1 h, *in vitro*) using AHL-PuP-2 (N7) as the detection probe. Volcano plots show the distribution of significantly liganded sites (competition ratio or CR<0.5, p<0.01) between the compound- and DMSO vehicle-treated groups in HEK293T soluble (**e**) and membrane proteome (**f**). The probe-modified peptides represent the combined results from n = 3 independent biological replicates. **g** and **h,** Competitive SuPUR TMT-ABPP analysis of ZH-2-055 and ZH-5-019 (25 µM, 1 h, *in vitro*) using ZH-2-087 (N9) as a probe. Significantly liganded sites (CR<0.5, p<0.01) between the compound- and DMSO vehicle-treated groups in HEK293T soluble (**g**) and membrane proteome (**h**) are depicted in volcano plots. The probe-modified peptides depicted are the combined results from n = 3 independent biological replicates. **i,** Protein-SuPUR ligand interaction map highlighting liganded sites (CR<0.5, p<0.01) from AHL-PuP-2 and ZH-2-087 competitive TMT-ABPP studies. Asterisk and pound symbols indicate a site located in a predicted druggable pocket by bioinformatics or structural modeling, respectively. Sites not detected in the respective TMT-ABPP studies (n.d.). The CR value represents an average across all peptide isoforms detected. **j**-**m,** Structural modeling using available crystal structures or predicted AlphaFold structures of SuPUR ligand bound to the active site of ABHD10 (Y215, AF-Q9NUJ1-F1, **j**) and NAD+ binding pocket of ABAD (Y168, PDB: 1U7T, **k**),^43^ catalytic domain of PARP2 (Y455, PDB: 8HLJ, pale green, **l**). Comparison of PARP2 with the homologous site on PARP1 (Y889, PDB: 7AAD, wheat, **l**) shows high degree of similarity in their substrate-binding pockets (**l**).^44, 45^ The liganded Y165 site on GNPNAT1 s positioned within the substrate catalytic pocket and forms hydrogen bonds with CoA, highlighting its role in enzymatic activity (PDB: 2O28, **m**).^46^ A yellow dashed lines represent hydrogen bond interactions. **n,** Competitive TMT-ABPP analysis of HEK293T proteome treated with ZH-1-049-2 (25 µM, *in vitro*, 1 h) or control ZH-2-036 (25 µM, *in vitro*, 1 h) versus DMSO vehicle revealed inhibition of ABHD10 and PPME-1 using FP-biotin activity-based probe (n = 2). **o,** The inhibitory activities of ZH-1-049-2, ZH-2-036, and the reported ABAD inhibitor Frentizole against ABAD were investigated using NAD/NADH-Glo biochemical assay. Data shown are mean ± SD; n = 3. **p,** ZH-1-049-2 exhibited selective liganding of PARP2 Y455, but no detectable binding activity at the corresponding PARP1 Y889 site as quantified by competitive TMT-ABPP (n = 3). **q,** ZH-1-049-2 exhibits selective inhibition of PARP2 activity while the PARP drug Olaparib shows equipotent inactivation of PARP1 and PARP2 as measured by a chemiluminescent biochemical assay. Data shown are mean ± SD. More examples are provided in Extended Data Fig. 3.

Next, we performed competitive TMT-ABPP deploying AHL-PuP-2 as the detection probe to identify ligandable sites and protein targets of ZH-1-049-2. In brief, HEK293T proteomes were pretreated with SuPUR ligand (25 µM, 1 h) followed by SuPUR probe labeling (100 µM, 1 h), CuAAC conjugation of desthiobiotin-azide, trypsin digestion, TMT labeling, avidin chromatography and LC-MS/MS analysis of probe-modified peptides (Extended Data Fig. 3a). These quantitative chemoproteomic studies demonstrated the SuPUR ligand ZH-1-049-2 but not the N-delete ZH-2-036 control engaged multiple target proteins with liganded sites found in predicted druggable pockets, including ABHD10 Y215, GSTP1 Y8, ABAD Y168, GNPNAT1 Y165, and PARP2 Y455 (CR<0.5, p<0.01; **Fig. 3e**, **f**). We confirmed these probe competition events were not due to protein expression changes (Extended Data Fig. 3b, c; Supplementary **Table 4**).

Subsequently, competitive TMT-ABPP was performed using the N9-SuPUR ligand ZH-2-055 and matching N9-ZH-2-087 probe. Chemoproteomic analyses revealed that ZH-2-055 but not the N-delete ZH-5-019 control retained covalent binding activity to several ZH-1-049-2 targets, including ABHD10, GSTP1, and ABAD (**Fig. 3g**, **h**). Akin to ZH-1-049-2 studies, these alterations from ZH-2-055 competition of probe binding were not due to protein expression changes (Extended Data Fig. 3d, e). By comparing ZH-1-049-2 and ZH-2-055 binding profiles, we identified several liganded sites in predicted drug pockets including PARP2 (Y455) and GNPNAT1 (Y165) that exhibited regioselective binding (**Figure 3i**, **l** and **m**).

### SuPUR ligand binds catalytic and non-catalytic sites to disrupt enzymatic functions

Several ZH-1-049-2-liganded sites are located within or near active sites prompting its further testing as a potential inhibitor of target proteins (**Fig. 3j**-**m**). We selected amyloid-β (Aβ) peptide-binding alcohol dehydrogenase (ABAD) and alpha/beta hydrolase domain-containing protein 10 (ABHD10) for initial evaluation. Covalent inhibitors of ABHD10, a serine hydrolase (SH) enzyme, are reported but inactivate principally through binding the catalytic and highly conserved active site serine residue.^47, 48^ Whether ABHD10 can be inactivated through binding of the non-catalytic tyrosine Y215 located near the catalytic serine (S152) is not known. We employed a gel-based competitive ABPP assay with the established activity-based probe fluorophosphonate-rhodamine (FP-Rh) to evaluate ZH-1-049-2 inhibitory activity against ABHD10.^49, 50^ The results indicated that ZH-1-049-2 exhibited moderate inhibitory activity, with an IC_50_ value of approximately 10 µM against a ∼34 kDa fluorescent protein band matching the expected MW of ABHD10 (Extended Data Fig. 3f). Next, we performed competitive ABPP using FP-biotin and a TMT-based MS readout to confirm target engagement against ABHD10. Quantitative ABPP analysis identified 65 SHs, with ZH-1-049-2 but not the N-delete ZH-2-036 control showing 40% inhibition of ABHD10 activity (25 µM compounds, **Fig. 3n** and Extended Data Fig. 3g).

Next, we tested whether covalent binding of ZH-1-049-2 to Y168 affects ABAD activity. ABAD is a mitochondrial dehydrogenase involved in steroid metabolism and reported to interact with intracellular amyloid-beta resulting in neuronal dysfunction in Alzheimer’s disease (AD).^51^ Currently there is a lack of potent and selective inhibitors that directly bind ABAD.^52^ Docking studies revealed that ZH-1-049-2 occupies the NAD+ binding pocket of ABAD,^43^ with the sulfonyl group forming hydrogen bonds (H-bonds) with Y168 and K172 (Extended Data Fig. 3h). To assess the biochemical inhibitory activity of ZH-1-049-2 and ZH-2-036, we established an NAD(P)/NAD(P)H-Glo assay for ABAD (see **Methods** for details). ZH-1-049-2 showed moderate inhibitory activity, with an IC_50_ of ∼1 µM (**Fig. 3o**). In contrast, the control compound ZH-2-036 exhibited weak inhibitory activity, while the reported ABAD inhibitor Frentizole^53, 54^ showed no activity even at the highest concentration tested (100 µM, **Fig. 3o**).

### SuPUR ligands inactivate PARP2 with high PARP isoform selectivity

The ability to achieve selectively within related members of an enzyme family is important for progressing fragment hits in ligand discovery programs. We were intrigued by the activity of ZH-1-049-2 against PARP2 but not PARP1 detected by chemoproteomics since ZH-1-049-2 binds the conserved Y455 site in the NAD+ binding pocket of the active site that is also engaged by the drug Olaparib (**Fig. 3l**, **p**; Extended Data Fig. 3i).^44^ A PARP biochemical assay was used to determine whether ZH-1-049-2 functions as a PARP2-selective biochemical inhibitor. Treatment with ZH-1-049-2 showed potent and concentration-dependent blockade of PARP2 (IC_50_ = 273 nM) and negligible activity against PARP1 even at the highest concentration tested (200 µM, **Fig. 3q**). As expected, treatment with Olaparib under the same experimental conditions resulted in equipotent inactivation of PARP1 and PARP2 (IC_50_ of ∼1 nM, **Fig. 3q**; see **Methods** for additional details of assay).

These data combined with findings from ABHD10 and ABAD support SuPUR ligands as inhibitors of diverse enzymatic functions with varying degrees of ligand efficiency (micro- to nano-molar potency) and in some cases with an unexpected degree of isoform selectivity.

### Disrupting catalytic activity of a serine hydrolase by liganding a non-catalytic tyrosine

The ability of ZH-1-049-2 to block diverse enzymes questioned whether this scaffold could be further optimized into a targeted covalent inhibitor. If demonstrated, the SuPUR chemotype could expedite fragment-based ligand discovery (FBLD) by reducing the screening library size needed for discovery campaigns. We elected to pursue ABHD10 for initial evaluation because this proof-of-concept target would establish a distinct inhibitory mechanism (i.e., binding to a non-catalytic tyrosine on SHs) for enabling ligand discovery against this important drug target class.^55, 56^

Molecular docking revealed that the terminal phenyl group of ZH-1-049-2 extends into a solvent-exposed pocket, suggesting that substitutions could be introduced at this position to improve binding affinity (Extended Data Fig. 4a). We synthesized and evaluated by competitive gel-based ABPP *in vitro* a focused library of 24 SuPUR ligands bearing modifications to the phenylsulfone region (2 µM compounds, 1 h; Extended Data Fig. 4b, c). At these lower concentrations, most compounds were inactive except for the 2,6-dichloro-substituted analog ZH-2-025 (**9**), which we confirmed to be a potent ABHD10 inhibitor in cells (IC_50_ of ∼400 nM, Extended Data Fig. 4d). Docking of ZH-2-025 into the ABHD10 active site provided key insights into the dramatic increase in potency compared with ZH-1-049-2; specifically, the chlorine atom of ZH-2-025 forms an important H-bond with arginine 280 (Extended Data Fig. 4e).

Next, we synthesized and screened by competitive gel-based ABPP a library of ZH-2-025 analogs (10 in total) and identified two candidates ZH-2-097 (-OCF3 substitution, **10**) and ZH-2-103 (-CF3 substitution, **11**) with improved potency (∼2-fold). The former was chosen as the optimized lead compound based on improved selectivity against a prominent ∼45 kDa off-target fluorescent protein band (Extended Data Fig. 4e, f). Direct comparison of ZH-2-097 with the initial hit compound ZH-1-049-2 showed a >300-fold enhancement in potency from a single modification to the parent ligand scaffold (**Fig. 4a**, **b**). Docking studies showed that ZH-2-097 could occupy the serine catalytic pocket, with the OCF_3_ substituent forming key H-bonds with Y87 (**Fig. 4c** and Extended Data Fig. 5a). The matching N9-isomer ZH-2-098 (**12**) also showed equipotent inhibitory activity against ABHD10 (**Fig. 4b**).

**Fig. 4.**
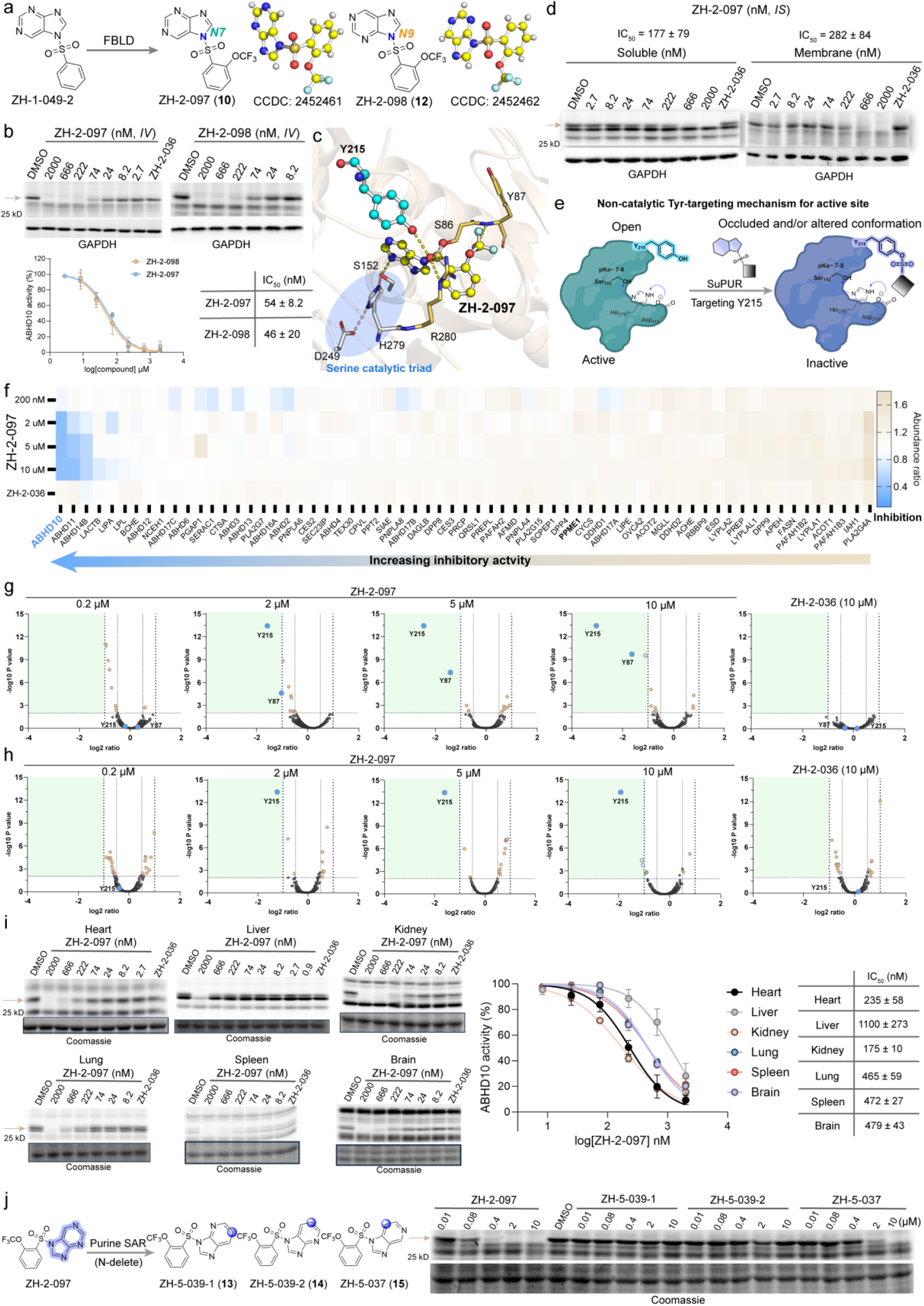
Disrupting the catalytic site of ABHD10 via SuPUR liganding of Y215. **a,** The potent N7-isomer ZH-2-097 (**10**) inhibitor was developed using fragment-based ligand discovery (FBLD). Additionally, the matching N9-isomer ZH-2-098 (**12**) was synthesized to evaluate regioselective activity of ligands. Crystal structures confirming regioisomeric state of ligand pairs are shown. **b,** Competitive gel-based ABPP analysis of ZH-2-097 and ZH-2-098 in HEK293T membrane proteomes using FP-Rh (*in vitro*, rt, 1 h). Data shown are mean ± SD (n = 3). **C,** Docking of ZH-2-097 into human ABHD10 (AF-Q9NUJ1-F1) supports ZH-2-097 binding in the serine catalytic pocket and hydrogen-bond interactions (yellow dashes) with Y215, Y87, S86, S152, and R280. Docking was performed with AutoDock Vina. The figure was generated using PyMOL 2.3.0. **d,** FP-Rh-mediated, competitive gel-based ABPP analysis of ZH-2-097 and control ZH-2-036 in proteomes from treated HEK293T cells (*in situ*, 2 h) in HEK293T proteome. Data shown are mean ± SD (n = 3). **e,** The proposed model of ABHD10 inhibition by ZH-2-097 involves covalent targeting of a non-catalytic tyrosine (Y215) to occlude the active site and/or alter protein conformation. **f,** FP-biotin-mediated Competitive TMT-ABPP analysis of HEK293T cells treated with ZH-2-097 (0.2-10 µM) and ZH-2-036 (control) versus DMSO vehicle revealed selective inhibition of ABHD10 across the SH superfamily. The FP-biotin-enriched SH proteins represent the combined results from n = 3 independent biological replicates. **g** and **h,** Competitive SuPUR TMT-ABPP of ZH-2-097 and ZH-2-036 in HEK293T cells. Volcano plots show concentration-dependent competition (CR<0.5, p<0.01) of ABHD10 Y215 and Y87 to a lesser degree compared with DMSO vehicle treated cells in HEK293T soluble (**g**) and membrane (**h**) proteomes. The AHL-PuP-2 modified peptides represent the combined results from n = 3 independent biological replicates. **i,** FP-Rh-mediated competitive gel-based ABPP of soluble proteomes from mouse tissues treated with ZH-2-097 indicated this ligand exhibited nanomolar inhibitory activity against ABHD10 detected in heart, liver, kidney, lung, spleen, and brain. Data shown are mean ± SD (n = 3). **j,** The binding activities of ZH-2-097 and corresponding nitrogen delete analogs (**13**-**15**) were assessed using competitive FP-Rh-mediate competitive gel-based ABPP in HEK293T soluble proteome. Data shown are mean ± SD (n = 3). More examples are provided in Extended Data Fig. 4 and 5. The N-delete analogs ZH-5-039-1 (**13**), ZH-5-039-2 (**14**), and ZH-5-037 (**15**) were synthesized to investigate the contribution of purine recognition to activity of ABHD10 inhibitors. Gel-based ABPP analysis indicated that only sulfonyl compounds with purine (ZH-2-097) exhibited nanomolar binding affinity for ABHD10, highlighting the purine backbone as a crucial recognition group (Fig. 4j). In summary, ABHD10 serves as a model example for developing potent and proteome-wide selective covalent inhibitors of SHs through SuPUR binding to a non-catalytic tyrosine.

We confirmed cellular activity of ZH-2-097 against ABHD10 by competitive gel-based ABPP analysis of proteomes from SuPUR ligand-treated HEK293T cells (IC_50_ ∼200 nM, **Fig. 4d**). To test our working hypothesis that SuPUR inhibitors of ABHD10 achieve selectivity through binding a non-catalytic tyrosine (Y215), we first assessed ZH-2-097 activity within the SH superfamily using FP-biotin.^49, 50^ Competitive TMT-ABPP analysis of proteomes from ZH-2-097-treated cells showed dose-dependent ABHD10 inactivation with remarkable SH-wide selectivity including removal of PPME1 off-target activity observed with the lead compound ZH-1-049-2 (**Fig. 4e**, **4f**, **3n**; Extended Data Fig. 5b-d).

To confirm tyrosine binding-mediated inactivation and proteome-wide selectivity, we subjected proteomes from ZH-2-097-treated cells to competitive TMT-ABPP using AHL-PuP-2 probe. Across 6,195 quantified binding sites, ZH-2-097 but not ZH-2-036 treatment resulted in significant and concentration-dependent liganding of ABHD10 Y215 (and Y87 to a lesser degree) with exquisite proteome-wide selectivity (CR<0.5, p<0.01; **Fig. 4g**, **h**). The observed competition events were not due to protein expression changes (Extended Data Fig. 5e). Finally, we showed that ZH-2-097 showed potent inhibitory activity against endogenous mouse ABHD10 detected across multiple tissues subjected to gel-based ABPP analysis (**Fig. 4i**).

### Proteome wide-selective inactivation of the multifunctional ABAD using regioselective SuPUR ligands

ABAD is a key enzyme involved in steroid metabolism, and its binding to Aβ is thought to contribute to neurotoxicity (Extended Data Fig. 6a). ABAD inactivation is a promising therapeutic strategy for the treatment of Alzheimer’s disease and various cancers.^57^ However, to date, only a limited number of small molecules targeting ABAD with moderate binding affinities have been reported.^52^ Here, we sought to test whether the lead compound ZH-1-049-2 could be rapidly developed into a potent ABAD inhibitor by targeting the catalytic Y168 site.

We deployed competitive gel-based ABPP to confirm ZH-1-049-2 activity is mediated through the Y168 site. Mutation of the catalytic tyrosine (Y168) largely abolished AHL-PuP-2 labeling of Y168G mutant compared with wild-type (WT) ABAD (**Fig. 5a**). We also confirmed the ABAD Y168G mutant showed a complete loss of biochemical activity, supporting SuPUR probe binding as a surrogate readout of catalytic activity (**Fig. 5b**). Gel-based competitive ABPP was performed to estimate potency of ZH-1-049-2 against recombinant ABAD-HEK293T proteome, yielding an IC_50_ value of approximately 5 µM (Extended Data Fig. 6b). We also found that ABAD prefers NAD+ over NADP+ as a cofactor for estradiol oxidation, matching previously reported findings (**Fig. 5b** and Extended Data Fig. 6c).^58^ Notably, our ABAD microplate-based biochemical assay exhibited a Z’ factor of ∼0.6, which indicates its suitability for high-throughput screening of ABAD inhibitors (Extended Data Fig. 6c).

**Fig. 5.**
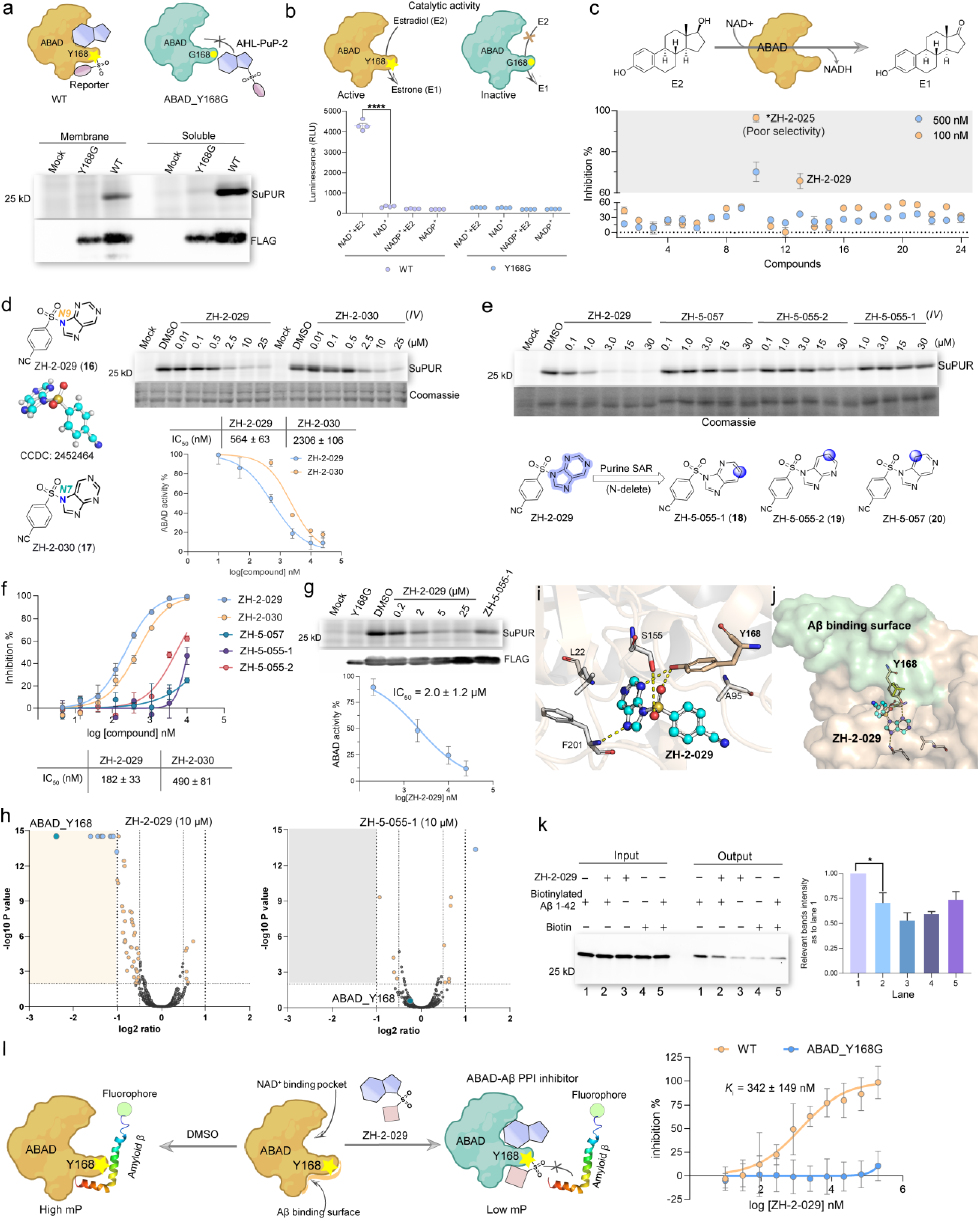
Disrupting catalytic and PPI activity of ABAD using a Y168-specific and N9-regioselective SuPUR ligand. **a,** Confirming enhanced nucleophilicity of catalytic Y168 by demonstrating loss of AHL-PuP-2 (25 µM) labeling of ABAD Y168G mutant compared with wild-type (WT) protein as measured by gel-based ABPP. **b,** Development of a NAD(P)/NAD(P)H-Glo assay to assess catalytic function of ABAD Y168 site and preference for NAD+ compared with NADP+ as cofactors using estradiol (E2) as the substrate. p values were analyzed by a one-way ANOVA comparing activity in presence versus absence of E2 (control group), ****p<0.0001) **c,** Screening of a SuPUR library (Supplementary Fig. 3) using the NAD(P)/NAD(P)H-Glo assay identified ZH-2-025 (**9**) and N9-isomer ZH-2-029 (**16**) as potent ligands against ABAD. Asterisk indicates ZH-2-025 exhibited poor selectivity toward ABAD because of potent ABHD10 off-target activity (Extended Data Fig. 6). **d,** Competitive gel-based ABPP (*in vitro*, 1 h) profiling shows enhanced potency for N9-isomer ZH-2-029 (**16**) compared to matching N7-isomer ZH-2-030 (**17**). Dose response (IC_50_) curves are shown as mean ± SD (n = 3). **e,** The nitrogen-delete analogs (**18**-**20**) showed impaired or no activity compared with parent ZH-2-029 (**16**) as determined by competitive gel-based ABPP. **f,** The ABAD inhibitory activity of SuPUR ligands and corresponding N-delete analogs were determined by NAD(P)/NAD(P)H-Glo assay. Data shown are mean ± SD (n = 3). **g,** Cellular potency of ZH-2-029 and matching N-delete control ZH-5-055-1 were assessed using gel-based competitive ABPP in HEK293T cells (*in situ*, 2 h). Data shown are mean ± SD (n = 3). **h,** Evaluation of target engagement and proteome-wide selectivity of ZH-2-029 and matching N-delete control ZH-5-055-1 in recombinant ABAD expressing HEK293T cells as determined by competitive TMT-ABPP using AHL-PuP-2 (*in situ*, 2 h). ZH-2-029 significantly ligands (CR<0.5, p<0.01) ABAD Y168 in cells in a site-specific and proteome-wide selective manner. The AHL-PuP-2 modified peptides represent the combined results from n = 3 independent biological replicates (n = 3). **I** and **j,** Docking of ZH-2-029 crystal structure into ABAD protein structure (PDB: 1U7T) revealed the ligand occupies the NAD+ pocket and forms H-bonds with S155, F201, and Y168 (**i**) and in proximity to the binding surface between ABAD and Aβ (**j**). Docking was performed with AutoDock Vina. The figure was generated using PyMOL 2.3.0. **k,** Pull-down assay was performed to demonstrate ZH-2-029 disrupts the protein-protein interaction (PPI) between ABAD and Aβ peptide. Date shown are mean ± SD, \**p*<0.05 (n = 3). **l,** Development of a fluorescence polarization (FP) binding assay to demonstrate dose–dependent blockade of fluorescent Aβ peptide binding to purified ABAD WT but not Y168G purified protein. Data shown are mean ± SD (n = 3). More examples are provided in Extended Data Fig. 6

We screened a library of ZH-1-049-2 analogs (24 in total) for hits displaying blockade of biochemical activity and identified ZH-2-025 and ZH-2-029 (**16**) as potent covalent ABAD inhibitors (IC_50_ of 60 and 180 nM, respectively; **Fig. 5c** and Extended Data Fig. 6d-e). While ZH-2-025 is more potent against ABAD, this ligand showed poor selectivity as evidenced by potent ABHD10 off-target activity (Extended Data Fig. 6f). Thus, ZH-2-029 was chosen for further evaluation. We confirmed by gel-based ABPP that the N9-isomer ZH-2-029 is a potent covalent ligand of ABAD (IC_50_ = 560 nM, **Fig. 5d**). The matching N7-isomer ZH-2-030 (**17**) was synthesized and found to exhibit a ∼3-4-fold reduction in potency against ABAD, supporting regioselective binding by ZH-2-029 (**Fig. 5d**).

The N-delete analogs ZH-5-055-1 (**18**), ZH-5-055-2 (**19**), and ZH-5-057 (**20**) were synthesized and tested to systematically evaluate the structure-activity relationship (SAR) of SuPUR ligand activity. Gel-based competitive ABPP results demonstrated that only the purine-containing ZH-2-029 exhibited nanomolar potency against ABAD, highlighting purine recognition as a key factor for inhibitory activity of ABAD ligands (**Fig. 5e**). Biochemical assay results further confirmed that only SuPUR ligands with a purine retained potent and regioselective inhibitory activity against ABAD, with ZH-2-029 and ZH-2-030 exhibiting IC_50_ values of 180 and 500 nM, respectively (**Fig. 5f**).

To test cellular activity of ZH-2-029, recombinant ABAD-expressing HEK293T cells were treated with ZH-2-029, followed by gel-based competitive ABPP analysis. The results indicated that ZH-2-029 but not the N-delete ZH-5-057 control covalently engages ABAD in living cells with moderate potency (IC_50_ = 2 µM, **Fig. 5g**). We performed competitive TMT-ABPP to confirm target engagement at the ABAD Y168 site and to assess proteome-wide selectivity. Evaluation of proteomes from ZH-2-029-but not ZH-5-055-1-treated recombinant ABAD-HEK293T cells revealed ABAD Y168 as the most significantly liganded site with evidence for proteome-wide selectivity across 6,000+ quantified binding sites (CR<0.05, p<0.01; **Fig. 5h** and Extended Data Fig. 6h). The observed competition events were not due to changes in ABAD protein expression (Extended Data Fig. 6g, i).

### SuPUR liganding of the catalytic Y168 site disrupts ABAD-Aβ PPI

ABAD is a multifunctional protein involved in regulation of mitochondrial metabolism, protein-protein interactions (PPIs) with intracellular Aβ,^51^ and tRNA processing as a component of the mitochondrial ribonuclease P complex.^59^ Molecular docking revealed that ZH-2-029 could form H-bonds with Y168, S155, and F201 in a binding site proximal to a PPI interface (highlighted in pale green) between ABAD and Aβ, suggesting that ZH-2-029 could disrupt the ABAD-Aβ interaction (**Fig. 5i**, **j**). Initially, we performed an affinity purification study to assess whether ZH-2-029 could disrupt enrichment of recombinant ABAD using a biotinylated Aβ peptide. The pull-down assay provided initial supporting evidence that the ABAD-Aβ interaction was blocked by ZH-2-029 (**Fig. 5k**).

Next, we established a fluorescence polarization (FP) assay to directly monitor purified ABAD interaction with a fluorophore-labeled Aβ peptide for assessing ABAD-Aβ PPI activity (Extended Data Fig. 6j). Interestingly, the ABAD Y168G mutant displayed impaired Aβ peptide binding affinity compared with WT protein (∼12-fold reduction in the calculated *K*_d_; Extended Data Fig. 6j). We detected concentration-dependent blockade of ABAD-Aβ interaction using ZH-2-029 with a calculated binding affinity matching potency values observed in the biochemical assay (*K_i_* = 342 nM, **Fig. 5l**).

In summary, we discovered that ABAD Y168 is a ligandable site for perturbing both catalytic and PPI activity using SuPUR ligands that regioselectively engage this catalytic residue with demonstrated proteome-wide selectivity.

### Phenotypic discovery of a site-specific, regioselective SuPUR inhibitor of ACAT2

We conducted disease ontology (DO) enrichment analysis to determine whether protein classes associated with disease pathology were overrepresented in our SuPUR interactome dataset (**Fig. 6a**).^60^ This analysis revealed significant enrichment of proteins involved in cancer cell proliferation, especially for gastrointestinal (GI) cancer and squamous cell carcinoma (SCC). Guided by these findings, we pursued phenotypic screening of our SuPUR ligand library to identify compounds that could block cell proliferation in a cancer cell type-specific manner. Subsequent SuPUR chemoproteomics would expedite target identification to reveal ligands for known and unexpected dependencies.

**Fig. 6.**
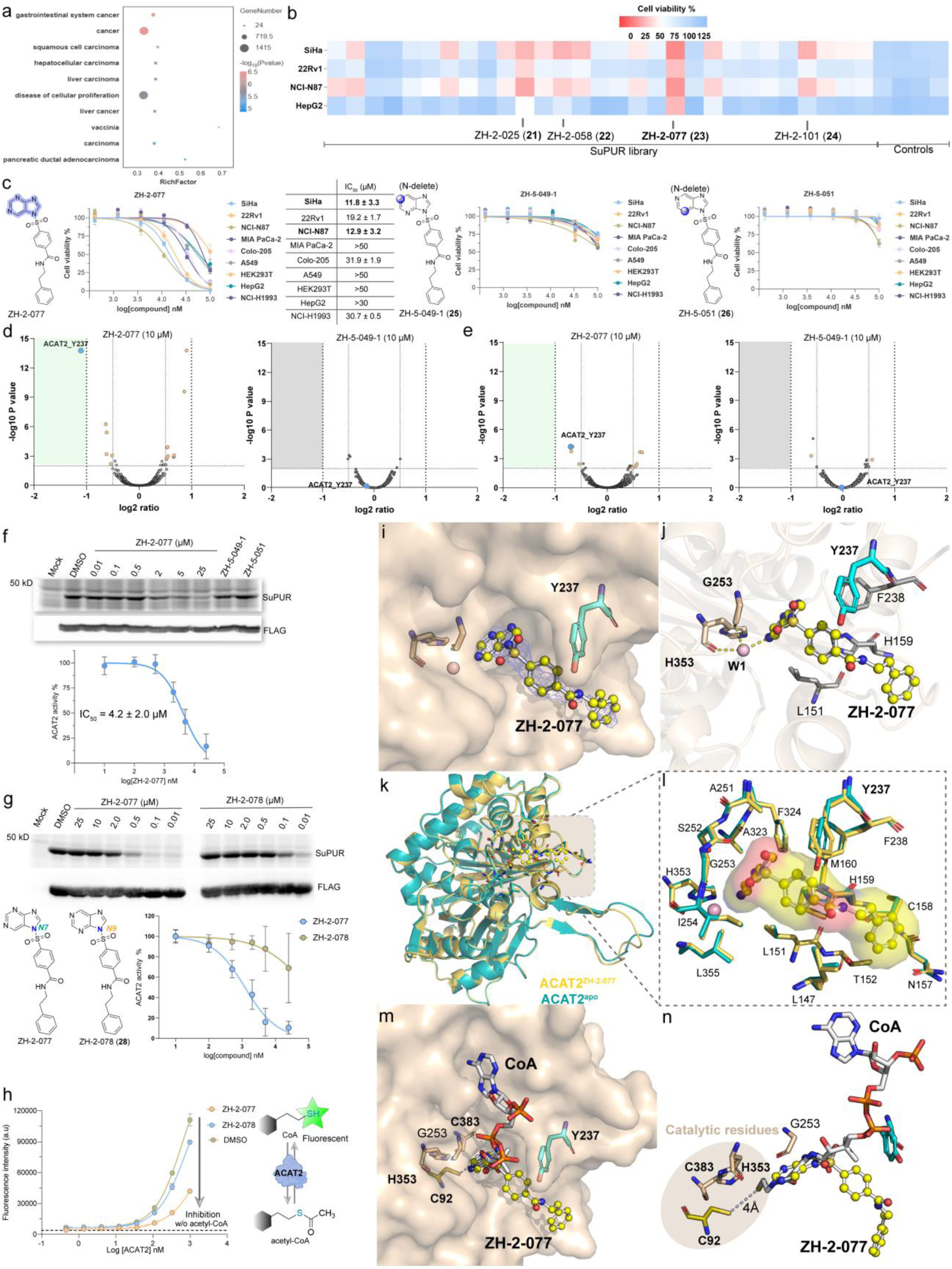
Targeted anti-cancer activity via a Y237-specific and N7-regioselective SuPUR inhibitor of ACAT2. **a,** Disease ontology (DO) enrichment analysis (GENE DENOVO^60^) was performed using protein detected in the collective AHL-PuP-2 dataset. **b,** Phenotypic screening of a SuPUR ligand library (30 µM, n = 2; Supplementary Fig. 4) to identify anticancer agents using a luminescent cell viability assay. **c,** Potency (IC_50_ values) of ZH-2-077 (**23**) and matching nitrogen-delete analogs ZH-5-049-1 (**25**) and ZH-5-051 (**26**) were evaluated using a luminescent cell viability assay across multiple cancer cell lines and noncancerous HEK293T cells. Data shown are mean ± SD (n = 3). **d** and **e,** Competitive SuPUR TMT-ABPP evaluation of target engagement and proteome-wide selectivity of ZH-2-077 and nitrogen-delete ZH-5-049-1 control in treated SiHa cells (*in situ,* 2 h). ZH-2-077 treatment results in significant liganding (CR<0.5. p<0.01) of ACAT2 Y237 in a site-specific and proteome-wide selective manner in SiHa soluble (**d**) but not membrane proteome (**e**). ACAT2 is expressed principally in the soluble fraction (data not shown). ZH-5-049-1 was inactive in both fractions as expected. The AHL-PuP-2 modified peptides represent the combined results from n = 2 independent biological replicates. **f,** Cellular potency of ZH-2-077 and matching nitrogen-delete controls ZH-5-049-1 and ZH-5-051 in recombinant ACAT2 expressing HEK293T cells as measured by gel-based competitive ABPP analysis (*in situ*, 2 h). Data shown are mean ± SD (n = 3). **g,** Comparing the N7-isomer ZH-2-077 and matching N9-isomer ZH-2-078 (**28**) by competitive gel-based ABPP in recombinant ACAT2-HEK293T soluble proteomes shows N7-regioselective binding of ACAT2 (*in vitro,* 1 h). Data shown are mean ± SD (n = 3). **h,** Biochemical assay shows regioselective inactivation of ACAT-mediated production of CoA by ZH-2-077 (50 µM). Titrations of purified ACAT2 were used for analyses. CoA levels were quantified using CoA Green, which detects the thiol (–SH) group in CoA by fluorescence. Data shown are mean ± SD (n = 3). **i,** View of the ACAT2^ZH-2-0^^77^ complex with the 2*mF_o_* – *DF_c_* electron density map for ZH-2-077 contoured at 1.0 σ with blue color. ACAT2: wheat; ZH-2-077: yellow sticks (PDB ID: 9V13, 2.5 Å). **j,** Binding mode of ZH-2-077 with ACAT2 (PDB ID: 9V13). The key interactions were highlighted as yellow dashed lines. **k,** Structural superimposition of the ACAT2 apo form (PDB ID: 9V37, 2.2 Å) with the ZH-2-077 bound ACAT2 complex (PDB ID: 9V13). **l,** Residues within a 5 Å radius of the ligand ZH-2-077 were highlighted to characterize the local binding environment. **m,** Superimposition of a ACAT2^ZH-2-077^ structure with ACAT2 complexed with CoA (PDB ID: 1WL4, white);^65^ the purine core of ZH-2-077 could occupy the catalytic pocket typically engaged by CoA. **n,** Highlighting the key catalytic residues, including the three catalytic residues C92, H353, and C383, with C92 playing a central role in its acetyltransferase activity. Additional examples are provided in Extended Data Fig. 7 and 8.

A panel of tumor cell lines was treated with SuPUR ligands along with N-delete controls (30 µM compounds) and cell viability was measured after 3 days of treatment. Several SuPUR ligands exhibited greater than 60% blockade of SiHa cell proliferation (compounds **21**-**24**; **Fig. 6b** and Extended Data Fig. 7a). Subsequently, we compared potency (IC_50_) for this subset of compounds in SiHa cells and identified ZH-2-077 (**23**) as the most active hit compound (Extended Data Fig. 7a). N-delete analogs of ZH-2-077 were synthesized, tested and found to lack inhibitory effects on cell proliferation, supporting the importance of purine recognition underlying ZH-2-077 activity (compounds **25**-**27**; **Fig. 6c** and Extended Data Fig. 7b). The cell viability results showed cell type-specific killing by ZH-2-077; this SuPUR ligand exhibited good inhibitory activity in SiHa and NCI-N87 cancer cells, with IC_50_ values of 12 and 13 µM, respectively (**Fig. 6c**). Importantly, no significant inhibitory activity was observed for these compounds in the non-cancerous HEK293T cell line.

Next, competitive TMT-ABPP was performed to identify the target protein(s) and binding site(s) engaged by ZH-2-077 in SiHa cells. Remarkably, across the 5,785 quantified Y and K sites, we identified Y237 on acetyl-CoA acetyltransferase 2 (ACAT2, UniProt ID: Q9BWD1) – a key enzyme involved in fatty acid and ketone body metabolism – as the most significantly liganded site with evidence for proteome-wide selectivity (CR ∼0.5, **Fig. 6d**, **e**).^61, 62^ The matching N-delete control ZH-5-049-1 showed negligible activity under the same treatment conditions. The detected changes in probe competition were not due to protein expression changes (Extended Data Fig. 7c). We also performed competitive TMT-ABPP analyses in 22Rv1 prostate cancer cells, which showed sensitivity to ZH-2-077 blockade of proliferation (IC_50_ = 20 µM, Extended Data Fig. 7d). Akin to SiHa cells, ZH-2-077 covalently engaged ACAT2 Y237 with high proteome-wide selectivity, supporting ACAT2 as the principal target of this SuPUR ligand in treated cancer cell lines (**Fig. 6d**-**e**; Extended Data Fig. 7d-e).

Bioinformatics analysis of available clinical data and published studies indicated that ACAT2 is highly expressed in certain cancers and is closely associated with poor overall survival, particularly in squamous, prostate and gastric cancers,^63, 64^ which matched the cell type specificity profile of ZH-2-077 (Extended Data Fig. 7f). We performed further biochemical verification by mutating the Y237 site and demonstrating loss of AHL-PuP-2 probe binding in ACAT2 Y237F mutant compared with WT protein (Extended Data Fig. 7g). Furthermore, concentration-dependent blockade of recombinant ACAT2 probe labeling in lysates and live cells was observed with ZH-2-077 but not the matching N-delete controls as determined by gel-based competitive ABPP (**Fig. 6f** and Extended Data Fig. 7h).

To investigate whether ACAT2 blockade by SuPUR ligands was regioselective, we synthesized and tested the matching N9-isomer ZH-2-078 (**28**). Strikingly, competitive gel-based ABPP revealed ZH-2-078 was largely inactive against recombinant ACAT2 (**Fig. 6g**). We also confirmed regioselective recognition of ACAT2 with direct SuPUR probe labeling; AHL-PuP-2 (N7) showed more prominent labeling of recombinant ACAT2 compared with ZH-2-087 (N9) despite equivalent protein expression (Extended Data Fig. 7i).

We next asked whether these liganding events at Y237 translated into blockade of ACAT2 biochemical activity. Here, we developed a biochemical assay using purified ACAT2 and monitored the conversion of acetyl-CoA to produce free CoA that is subsequently quantified by fluorescence detection (Extended Data Fig. 7j). Using this assay, we confirmed regioselective blockade of ACAT2 biochemical activity using ZH-2-077 compared with ZH-2-078 and vehicle control (**Fig. 6h**).

### Crystal structure of ACAT2 bound to ZH-2-077

To gain further insights into ZH-2-077 binding mechanism, we crystallized ACAT2 in the absence (ACAT2^apo^) and presence of ligand (ACAT2^ZH-2-077^) and determined structures at 2.2 and 2.5 Å resolution, respectively (Supplementary **Table 5**). The ACAT2^ZH-2-077^ co-crystal structure was determined by molecular replacement using a previously reported ACAT2^CoA^ structure as the search model (PDB: 1WL4).^65^ Two polypeptide chains were identified and built within the asymmetric unit. Compared to ACAT2^apo^, the ACAT2^ZH-2-077^ complex structure revealed a distinct additional electron density in the CoA-binding site (Extended Data Fig. 8a, b). Subsequently, ZH-2-077 was modeled into the observed ligand density, allowing analysis of its binding environment and providing structural insights into ACAT2 inhibitory mechanism (**Fig. 6i**). Notably, a key water molecule, coordinated by the main-chain atoms of G253 and H353, exhibited an interaction with the N7 position of the purine group of ZH-2-077 (**Fig. 6j**).

Superimposition of the ACAT2^apo^ and ACAT2^ZH-2-077^ structures revealed minor conformational changes in loop regions and the ZH-2-077 binding pocket (surrounding residues within 5 Å, **Fig. 6k**, **6l**). Overlay of ACAT2^ZH-2-077^ complex with ACAT2^CoA^ revealed that the purine core of ZH-2-077 is positioned deep into the catalytic pocket (**Fig. 6m**, **6n**). Importantly, the bridging water molecule detected in the presence of ZH-2-077 was absent in the ACAT2^CoA^ structure (**Fig. 6j**). The SuPUR-liganded residue Y237 is located adjacent to the CoA-binding site suggesting that adduction here with ZH-2-077 would disrupt CoA interaction. The positioning of the ZH-2-077 purine in proximity to the conserved catalytic triad (C383, H353, and C92) provides further insights into how ligand binding disrupts CoA recognition in the ACAT2 active site (**Fig. 6k-n** and Extended Data Fig. 8c).

In summary, we discovered the SuPUR ligand ZH-2-077 as a first-in-class covalent ligand of ACAT2 and provide atomic level resolution of its inhibitory mechanism via regioselective binding to Y237. Notably, the proteome-wide selectivity of ZH-2-077 highlights the ability of SuPUR chemoproteomics to expedite phenotypic screening programs.

## Discussion

Purines are essential and structurally related bioactive molecules that interact with a large fraction of the human proteome (∼20%) to mediate key cell biology.^8^ The increasing number of drugs containing a purine core supports its key role in facilitating molecular recognition at functional protein pockets (**Fig. 1**). While certain protein-purine interactions (e.g., kinase-ATP recognition) have been extensively explored and targeted with reasonable specificity^66^, a global assessment of ligandable purine-binding pockets in the human proteome is largely underexplored.

We developed SuPUR chemoproteomics as an enabling framework to functionally interrogate the human purine interactome. These efforts produced a massive resource of 31,000+ targetable Y and K sites on proteins that showed substantial overlap (41%) with the reported purinome (**Fig. 2**). The SuPUR interactome represents, to the best of our knowledge, the most comprehensive chemoproteomic database for covalent targeting of residues beyond cysteine. To assess ligandability of purine binding pockets in the human proteome, we evaluated SuPUR ligands in lysates, cells and purified protein that collectively represent quantification of ∼1M data points that we provide in this report as a resource for the scientific community. A key discovery from these collective studies was the privileged nature of the purine scaffold for building proteome-wide selective protein ligands de novo (**Fig. 4**-**6**).

Regioselective modification on the N7 or N9 positions on purine is a mechanism for controlling biological specificity in cells (e.g., mRNA stability vs nucleoside biosynthesis, respectively).^23, 25^ We surmised that strategic placement of a sulfonyl electrophile at the N7 or N9 position will produce matching probes to facilitate regioselective ABPP of the SuPUR interactome. This approach is analogous to incorporation of stereocenters into electrophilic ligands for evaluating stereoselective binding, which helps discern more specific interactions vs general labeling of protein residues.^67–69^ Regioselective recognition to guide covalent ligand discovery, in contrast, has been largely unexplored and represents a distinct feature enabled by the SuPUR scaffold. In support of this hypothesis, sites liganded in a regioselective fashion displayed high isoform selectivity (PARP2) or could be rapidly progressed into proteome-wide selective inhibitors (ABAD and ACAT2, **Fig. 3**, **5** and **6**).

SuPUR chemoproteomics could be deployed for both target- and phenotypic-based discovery campaigns to produce potent ligands for functional pockets that inactivate proteins through distinct mechanisms. A surprising and recurrent theme from these programs was the minimal structural modifications to the SuPUR ligand scaffold required to achieve potency and proteome-wide selectivity (**Fig. 4**-**6**). These findings were initially counter-intuitive especially when one considers SuPUR probes – developed using the same chemotype – captured thousands of distinct binding sites by chemoproteomics (**Fig. 1** and **2**). A potential explanation is that SuPUR chemistry akin to reported sulfone-based electrophiles are highly tunable in protein reactivity and affinity.^29^ It is tempting to speculate the latter is more pronounced with the SuPUR scaffold because of molecular recognition imparted by the purine group; additional follow-up studies are needed to fully test this hypothesis.

We discovered unexpected inhibitory mechanisms using SuPUR targeting. The ABHD10 inhibitor (ZH-2-097) represents a proof-of-concept target for liganding a non-catalytic tyrosine to inactivate a SH with proteome-wide selectivity (**Fig. 4**). Members of this enzyme superfamily are important therapeutic targets that are typically inactivated by electrophilic probes and drugs via binding the conserved active site serine.^56, 70^ Our findings present a distinct inhibitory mechanism for SHs via covalent binding of a non-catalytic and less conserved tyrosine that results in either occlusion of the active site and/or alteration in protein conformation (**Fig. 4**). We also developed SuPUR ligands that disrupt the ABAD-Aβ PPI activity through liganding of the catalytic Y168 site (**Fig. 5**). The ability to disrupt both catalytic and non-catalytic functions of ABAD using a single agent has not been reported, to the best of our knowledge, and important given the multifunctional role of this enzyme in mitochondrial metabolism, tRNA processing and Aβ interactions.^51, 59^ Given the genetic evidence linking ABAD to human disease, we envision ZH-2-029 will be an important tool for enabling mechanistic studies on ABAD biology and pathology.

Phenotypic screening of a SuPUR ligand library for anticancer activity produced a site-specific (Y237), regioselective (N7) ACAT2 ligand displaying proteome-wide selectivity in cells (**Fig. 6**). These studies were significant for several reasons: (i) ZH-2-077 represents, to the best of our knowledge, the first ACAT2 inhibitor, (ii) the SuPUR platform can accelerate target identification and lead optimization in phenotypic screening campaigns, and (iii) ACAT2 is a metabolic dependency in gastrointestinal and squamous cell carcinoma cells. We determined the structures of apo- and ZH-2-077-bound ACAT2 to gain mechanistic insights into inhibition with atomic resolution. These structural studies revealed the importance of the N7-purine group of ZH-2-077 in forming key interactions with catalytic residues deep in the CoA binding pocket to align the electrophile in proximity to the adjacent Y237 site. Notably, a bridging water molecule that H-bonds with the N7-purine in the CoA pocket of ACAT2^ZH-2-077^ would likely be lost with a corresponding N9-purine in the same binding mode (**Fig. 6**). We speculate this interaction contributes to the exquisite regioselectivity for ZH-2-077-mediated ACAT2 inactivation and plan to investigate further in future studies.

Several areas for future inquiry are warranted to further advance the SuPUR platform. First, SuPUR chemistry in the current study is limited to the basic purine as a leaving group. We envision that deployment of modified purines (e.g., adenine or guanine) will alter the proteomic binding profiles and potentially capture a larger fraction of the purinome.^71^ Further studies are needed to understand whether prominent SuPUR binding to ABHD10 is indicative of a potential role in purine recognition or metabolism, which is not currently known for this reported depalmitoylase.^55^ Whether SuPUR liganding of ABAD affects its role in mitochondrial ribonuclease P activity would provide further insights into the catalytic and non-catalytic roles of Y168. While the SuPUR ligands for ACAT2 are highly selective, additional optimization efforts are needed to improve potency for potential applications *in vivo*.

In summary, we develop a chemoproteomic platform that enables regioselective targeting of the human purine interactome and provide multiple examples of its application for proteome-wide selective modulation of enzymatic and PPI function to expand the scope of intractable proteins addressable with small molecules.

## Supporting information

ACAT2_apo_PDB_validation

ACAT2_compound_PDB_validation

## Methods

### Expression and purification of ABAD_Y168G

Target DNA sequence encoding ABAD Y168G mutant was optimized and synthesized. The synthetic gene was subsequently cloned into vector pET30a with an N-terminal His tag for protein expression in *E. coli.* The recombinant plasmid was transformed into E. coli BL21 Star™ (DE3) cells. A single colony was selected and inoculated into LB medium supplemented with the appropriate antibiotic, followed by incubation at 37 °C with shaking at 200 rpm. Protein expression was induced by the addition of IPTG, and expression levels were monitored by SDS-PAGE. Recombinant BL21 Star™ (DE3) cells stored in glycerol were inoculated into TB medium supplemented with appropriate antibiotics and cultured at 37 °C. When the optical density OD_600_ reached about 1.2, protein expression was induced by IPTG at 15°C for 16 h. Cells were then harvested by centrifugation. Cell pellets were resuspended with lysis buffer and lysed by sonication. The supernatant after centrifugation was retained for protein purification. ABAD Y168G was purified by one-step purification using a Ni^2+^-NTA column. The protein was sterilized by 0.22 μm filter before being stored in aliquots. Protein concentration was determined using the Bradford protein assay with BSA as a standard. The protein purity and molecular weight were assessed by standard SDS-PAGE. We thank GenScript for protein expression and purification (Lot# U079U921G0).

### Expression and purification of ACAT2

The DNA fragment encoding ACAT2 (Accession: Q9BWD1) was synthesized and subcloned into the pET28a vector between the NcoI and XhoI restriction sites. The resulting plasmid was transformed into *Escherichia coli* BL21 (DE3). 2 L of LB medium was inoculated with 5 mL of an overnight culture grown in LB containing 50 μg/mL kanamycin. The culture was incubated at 37 °C until the OD_600_ reached 0.6 and followed by induction with 500 μL of 1 M isopropyl β-D-thiogalactopyranoside (IPTG). After 12 h induction at 16 °C, cells were harvested by centrifugation and resuspended in 30 mL of lysis buffer (20 mM Tris-HCl, pH 8.0, and 500 mM NaCl) supplemented with 1% Triton X-100. Cells were lysed by sonication, and the cell debris was removed by centrifugation at 13,000 × g. The supernatant was applied to a Ni-NTA affinity column, washed with 50 mL of lysis buffer. The recombinant ACAT2 with C-terminal His-tag was eluted with 25 mL of elution buffer (20 mM Tris-HCl, pH 8.0, 500 mM NaCl, and 1 M imidazole). The eluted protein was further purified by size-exclusion chromatography using a HiLoad 16/60 Superdex 200 pg column, equilibrated with buffer containing 20 mM Tris-HCl, pH 7.6, and 200 mM NaCl. Protein purity was assessed by 10% SDS-PAGE, and protein concentrations were determined based on absorbance at 280 nm using the calculated molar extinction coefficient (ε_280_ = 26,970 M^−1^ cm^−1^).

### Crystallization and data collection

ACAT2 after protein purification was concentrated to 16 mg/mL in buffer containing 20 mM Tris-HCl, pH 7.6, and 200 mM NaCl, and crystallized using the sitting-drop vapor-diffusion method at 20 °C. Crystals were obtained using a reservoir solution composed of 10% (v/v) 2-propanol, 0.1 M BICINE, pH 8.5, and 30% (w/v) polyethylene glycol 1500. The complex structure ACAT2^ZH-2-077^ was generated by soaking the ACAT2 crystals with 0.5 mL of 5 mM compound ZH-2-077 in the crystallized condition for 15 min. Crystals were cryoprotected and flash-frozen in liquid nitrogen for data collection. X-ray diffraction data were collected at the National Synchrotron Radiation Research Center (NSRRC), Taiwan, on beamline 15A using 0.97 Å with an EIGER 2X CdTe 1M detector. Data was indexed and scaled using HKL2000 software.^72^

### Structure determination and refinement

The structure of ACAT2 bound ZH-2-077 (ACAT2^ZH-2-0^^77^) and apo-form (ACAT2^apo^) were solved by the molecular replacement (MR) method by MOLREP,^73^ using the structure of ACAT2 (PDB entry: 1WL4) as the search model.^65^ The Initial model was built using COOT.^74^ Structure refinement was performed with REFMAC.^75^ The final atomic coordinates and structural factors of the complex structure ACAT2^ZH-2-0^^77^ and ACAT2^apo^ have been verified and deposited in the Protein Data Bank (PDB) with the accession code 9V13 and 9V37, respectively.

### NAD(P)/NAD(P)H Glo assay

Assay buffer: 20 mM Tris pH7.4 with 0.002% Tween-20 and 0.02% BSA.

Substrate: 17β-Estradiol, 100 mM in DMSO.

Cofactor: NAD+ Grade I free acid, 20 mM in H_2_O.

ABAD purified protein (5 µL, 80 nM) was added to each 384 wells plate along with 2.5 µL of 4X compounds solution. Assay plate was incubated at room temperature (RT) for 60 min, and then 2.5 µL of 8X substrate/8X cofactor mix was added to each well for a final concentration of 50 µM estradiol and 500 µM NAD^+^. The assay plate was incubated at 37 °C for 3 h. NAD(P)H-Glo™ Detection System reagents were prepared according to manufacturer’s specifications, and 10 µL was added to each well. After incubating for 1 h at RT, luminescence (Lum) was measured using Agilent Bio-Tek multimode reader. The percentage inhibition was calculated using Equation 1 where Lum_blank_ is the signal for the free 17β-Estradiol (blank control) and Lum_DMSO_ is the signal for the DMSO treatment (no inhibition activity). And Lum is the readout value of compound treatment groups.

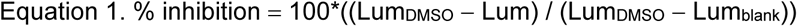

Plot dose-response curves and determines IC_50_ values using GraphPad Prism 10.0.

### Pull-down assay

Assay buffer: 10 mM Tris, 0.1 M NaCl, 1 mM EDTA.

Substrate: Biotin-β-Amyloid (1-42, Biotin-DAEFRHDSGYEVHHQKLVFFAEDVGSNKGAIIGLM VGGVVIA), 500 µM in 1%NH_4_OH.

The concentration of HEK293T cell lysate overexpressing ABAD was determined by DC (detergent compatible) protein assay. Following quantification, the biotin Aβ (1-42) was added to the 50 µL cell lysate (2 mg/mL) for a final concentration of 1 µM, and the mixture was incubated overnight at 4 °C. Add the immune system was then incubated with avidin agarose resin for 1 h at RT. To remove unbound proteins, the avidin-bound complex was washed 5 times with PBS (1400g for 1 minute). After that, the washed resin was resuspended in 30 µL PBS, followed by the addition of 4X loading buffer (10 µL per aliquot). The mixture was then boiled, centrifuged at 500g for 1 minute, and the resulting supernatant was used for electrophoretic analysis.

### Gel-based ABPP (*in vitro*)

HEK293T cell proteome were diluted to a final concentration of 1 mg/mL and aliquoted according to the required reaction conditions (49 µL for probe only and 48 µL for ligand competition assay). For competition experiments, proteomes were pretreated with 1 µL of 50X compound DMSO stock and incubated at RT for 1 h. Subsequently, 1 µL of 50X probe (1.25 mM) DMSO stock was added to each tube for a final concentration of 25 µM and incubated for 1 h at RT. After incubation, 6 µL of the click chemistry reagents were added to the mixture, consisting of: 1 µL Rh-N₃ (1.25 mM), 1 µL TCEP (50 mM), 3 µL TBTA (1.7 mM), 1 µL CuSO₄ (50 mM).

The reaction mixture was incubated at RT for 1 h to facilitate the click reaction. Finally, click reaction was quenched by adding 17 µL of 4X SDS-PAGE loading buffer with β-mercaptoethanol (βME), followed by vertexing. And the 30 µL of each sample was analyzed by SDS-PAGE (300V, 800Vh). The results were visualized using Bio-Rad ChemiDoc MP imaging system. The percentage of control was calculated using Equation 2 where FL_blank_ is the signal of blank group (blank control) and FL_DMSO_ is the signal for the DMSO treatment (no inhibition activity). And FL is the readout value of compound treatment groups.

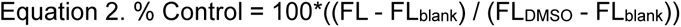

Plot dose-response curves and determines IC_50_ values using GraphPad Prism 10.0.

### Gel-based ABPP (*in situ*)

HEK293T living cell (∼1×10^6^ cells/well) treated with probes or ligands for 2 h in a 6-well plate with serum-free media. Subsequently, cells were harvested and washed 3 times using DPBS to remove unreacted compound or probe. Cells were lysed by sonication, and the resulting lysate was subjected to ultracentrifugation at 100,000 × g for 45 min to separate the soluble (supernatant) and membrane-associated (pellet) proteomes. HEK293T cell proteomes were diluted to a final concentration of 1 mg/mL and aliquoted according to the required volumes (50 µL for probe only and 49 µL for ligand competition). The remaining steps followed the gel-based ABPP (*in vitro*) protocol.

### Gel-based competitive ABPP with FP-Rh

HEK293T cell proteome were diluted to a final concentration of 1 mg/mL and aliquoted according to the required reaction conditions (49 µL for probe only and 48 µL for ligand competition assay). For competition experiments, proteomes were pretreated with 1 µL of 50X compound DMSO stock and incubated at RT for 1 h. Subsequently, 1 µL of 50X FP-Rh (50 µM) DMSO stock was added to each tube for a final concentration of 1 µM and incubated for 1 h at RT. Subsequently, the reaction was quenched by adding 12.5 µL of 4X SDS-PAGE loading buffer with β-mercaptoethanol (βME), followed by vertexing. And 30 µL of each sample was analyzed by SDS-PAGE (300V, 800Vh). The results were visualized using Bio-Rad ChemiDoc MP imaging system. The percentage of control was calculated using **Equation 2**, where FL_blank_ is the signal of blank group (blank control) and FLDMSO is the signal for the DMSO treatment (no inhibition activity). And FL is the readout value of compound treatment groups.

### Gel-based ABPP (tissue labelling)

C57BL/6J mice (male) were purchased from Jackson Laboratory (Ref# O-000007447). The mouse was sacrificed following the protocol AUP-2023-0026. And the tissues were collected for analysis. Approximately 30 mg of tissues were weighed and suspended in 1.0 mL of DPBS in 1.5 mL snap-cap tubes. The tissue was then homogenized using a Bullet Blender homogenizer. The mixture was then centrifuged at 3,000 × g for 3 min, and the protein concentration in the resulting supernatant was quantified using the BCA assay. The remaining steps followed the Gel-based competitive ABPP with FP-Rh.

### Western Blot (WB)

The HEK293T proteome was diluted to a final concentration of 1 mg/mL. 4X loading buffer containing β-mercaptoethanol was added to the samples. A total of 10 µL of each sample was loaded onto an SDS-PAGE gel, and electrophoresis was performed at 120 V for 55 min. Following electrophoresis, proteins were transferred onto a nitrocellulose membrane using the semi-dry transfer method at 15 V for 10 min. The membrane was then blocked with 5% BSA in TBST for 1 h at RT to prevent nonspecific binding. After blocking, the membrane was incubated with the primary antibody overnight at 4°C. The membrane was washed three times with TBST (10 mL) to remove unbound primary antibody. A fluorescent-conjugated secondary antibody was then added, and the membrane was incubated at RT for 1 h. Subsequently, the membrane was washed three times with TBST (10 mL), and protein bands were visualized using the ChemiDoc imaging system.

### Transfection

The transfection reagent mixture was prepared by combining 600 µL serum-free DMEM, 20 µL of polyethylenimine (PEI, 1 mg/mL, pH 7.4), and 2.6 µg of plasmid DNA. First, the mixture was incubated at RT for about 30 min to allow the formation of PEI-DNA complexes. HEK293T cells (∼1×10^6^) were cultured in a 100 mm cell plate with penicillin-streptomycin (PS) free cell media and transfected when cell confluency reached approximately 50%. The transfection mixture was then added dropwise to the cells, and the plate was gently swirled to ensure even distribution. The cells were incubated for 48 h to allow for target protein expression. Subsequently, the cell medium was aspirated, and the cells were washed two times with 10 mL DPBS. The cells were then craped into DPBS, collected in 10 mL conical tubes by centrifugation at 1000 x g for 3 min. Cells pellets were ready for Gel-ABPP analysis.

### PARP2/PARP1 chemiluminescent assay

The chemiluminescent assay was followed by Bioscience chemiluminescent assay kit protocol.

First, the plate was coated with 50 µL histone mixture at 4 °C overnight. The plate was washed three times with 200 µL PBST, then blocked with 200 µL blocking buffer and incubated at RT for 1.5 h. The plate was then washed three times with 200 µL PBST again. Immediately after, 50 nL of diluted compound was transferred from the compound stock plate into the assay plate using the Echo 550 liquid handler system. Subsequently, 30 μL of the master mixture (3 μL 10 × PARP buffer, 3 μL 10 × PARP assay mixture, 6 μL activated 5 × DNA, and 18 μL water) were added to each well. The enzyme-based reaction was initiated by adding 20 µL of PARP1 or PARP2 enzyme (2.25 ng/mL) to the wells. The blank control was designed by adding 20 μl of 1× PARP buffer only. The mixture was incubated at RT for 1 h and then discarded the reaction mixture. The plate was then washed three times with 200 μL PBST. 50 µl of diluted Streptavidin-HRP was added to each well and incubated for 30 min at RT. The mixture was discarded, and the plate was washed three times with 200 µL PBST buffer. Just before use, 50 µL of ELISA ECL substrate A and 50 µL of ELISA ECL substrate B were mixed on ice, and 100 µL of the mixture was added to each well. The chemiluminescence was measured by the Ensight reader system, The percentage inhibition was calculated using Equation 3 where Lum_blank_ is the signal for the free PARP2 enzyme (blank control) and Lum_DMSO_ is the signal for the no compound treatment (no inhibition activity). And Lum_Samples_ is the readout value of compound treatment groups.

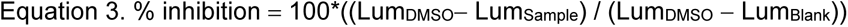

Plot dose-response curves and determines IC_50_ values using GraphPad Prism 10.0.

### Luminescent-based cell viability assay

The luminescent-based cell viability assay was followed by VaZyme CellCounting-Lite 2.0 kit protocol.

Human cells (5000 cells/well) were seeded into 96 transparent wells plate (Corning) with 50 µL of the appropriate cell medium, Cells were allowed to adhere overnight at 37 °C. Subsequently, Test compounds were prepared by diluting stock solutions to 2 × the final concentration in cell medium. And then 50 µL tested compounds were added to each well of the 96 wells plate. Cells were incubated with compounds for a period of 72 h. Next, the 96 well plate was taken out from the incubator and allowed to stand at RT for 30 min. Subsequently, cell counting-Lite 2.0 reagent (100 µL) was added to each well. The plate was gently shaken for 2 to 5 min to ensure thorough mixing and complete cell lysis. The mixture was left to stand at RT for 10 min to stabilize the luminescent signal. Then, the luminescent values were determined by Agilent Bio-Tek multimode reader.

The percentage cell viability was calculated using Equation 4 where Lum_blank_ is the signal for the free cells (blank control) and Lum_DMSO_ is the signal for the no compound treatment (0.1% DMSO). And Lum_Samples_ is the readout value of compound treatment groups.

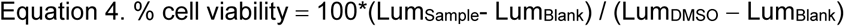

Plot dose-response curves and determines IC_50_ values using GraphPad Prism 10.0.

### Fluorescence polarization (FP) assay of ABAD-Aβ (1-42) binding^76^

FP assays were performed using a Synergy H4 Multi-Mode Microplate Reader (BioTek) with excitation 488/20 and emission 528/20. Fluorescence polarization experiments were performed in 384-well, flat bottom (6008260, ProxiPlate) in a final volume of 10 μL. The final reaction buffer contained: 4 nM fluorescent Aβ (1-42),10 mM tris-HCl, 150 mM NaCl, 0.01% Tween-20, pH 7.4. Dose-dependent experiments were performed using at least 8 concentrations of ABAD and ABAD_Y168G purified proteins in 3-fold serial dilutions from 3 µM and 30 µM, respectively. Subsequently, the reaction mixtures were incubated at RT for 60 min, then total fluorescence and fluorescence polarization measurements were taken. The binding affinity *K*_d_ value was determined from FP readouts using GraphPad Prism 10.0.

### Fluorescence polarization (FP) competition assay of ABAD

FP assays were performed using Synergy H4 Multi-Mode Microplate Reader (BioTek) with excitation and emission filters appropriate for the fluorophore used in the binding experiment. Fluorescence polarization experiments were performed in 384-well, flat-bottom, black assay plates (6008260, ProxiPlate) in a final volume of 10 μL. The final assay buffer contained: 10 mM tris-HCl, 150 mM NaCl, 0.01% Tween-20, pH 7.4, 0.01% Tween-20, and 1 µM ABAD or 10 µM ABAD_Y168G. The mixture was incubated for 60 min at RT, followed by the addition of 1 μL fluorescent Aβ (1-42), for the final concentration of 4 nM. Subsequently. The mixture was incubated for a minimum of 60 min at RT. Fluorescence polarization (FP) was measured from the top of the well with a Synergy H4 Multi-Mode Microplate Reader (BioTek) with polarized filters and optical modules for fluorescein (λex = 488 nm ± 20 nm, λem = 528 nm ± 20 nm). Dose-dependent experiments were performed using at least 10 concentrations of the test compounds in 3-fold serial dilutions starting from 200 μM. For each assay, negative controls (equivalent to 0% displacement) contained the fluorescent Aβ (1-42), ABAD or ABAD_Y168G, and 1 μL FP buffer; blank controls contained only the fluorescent Aβ (1-42) and 2 μL FP buffer. After 60 min of incubation at RT, the fluorescence polarization was measured using plate reader.

The percentage inhibition was calculated using Equation 5, where mP_free_ (blank) represents the signal for the free fluorescent Aβ (1-42) and mP_bound_ (negative) represents the signal of fluorescent Aβ (1-42) bound to target proteins.

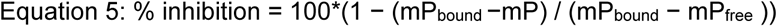

IC_50_ was determined for duplicate measurements by non-linear least-squares analysis using GraphPad Prism 10.0. The *K*_i_ value of the inhibitor was calculated using the Cheng-Prusoff equation.

Cheng-Prusoff equation: *K*_i_ = IC_50_ / (1 + [L]/*K*_d_) ^77^

The *K*_d_ value was determined using a constant concentration of Aβ (1-42) and titrating with ABAD protein at increasing concentrations. [L] represents the concentration of the ABAD protein used in the FP assay.

### ACAT2 biochemical assay

Coenzyme A (CoA) released from the acetyl-transfer reaction catalyzed by ACT2 was monitored using the fluorescent probe CoA Green (AAT Bioquest). Reactions were conducted in 384-well microplates (Corning, Cat#3575), and fluorescence was measured with excitation at 485 nm and emission at 535 nm. The reaction solution contained NaCl (137 mM), Na_2_HPO_4_ (8 mM), KH_2_PO_4_ (1.5 mM), KCl (2.7 mM), pH∼7.4. 10 µL of ACAT2 protein (100 nM) was added to 384 wells and followed by 10 µL of ACAT2 substrate acetyl-CoA (200 µM). The mixture was gently mixed and incubated at 37 °C for 3 h. Subsequently, 20 µL CoA green detection reagent was added, and the mixture was incubated at RT for 40 min. Fluorescence intensity was then recorded using a microplate reader (Ex/Em = 485/535 nm).

### ZH-2-077 and ZH-2-078 inhibitory activity against ACAT2

To evaluate compound activity, 5 µL of tested compounds (2×,100 µM) or 1% DMSO control was added to 384-well microplates. A dose titration of ACAT2 protein (2 ×) from 900 pM to 2000 nM in threefold dilutions was then added. The mixture was incubated at RT for 1 h, followed by the addition of 10 µL ACAT2 substrate acetyl-CoA (2×, 200 µM). Reactions were incubated at 37 °C for 3 h. Subsequently, 20 µL of CoA green detection reagent was added, and the plate was incubated at RT for an additional 40 min. Fluorescence was measured using a microplate reader (Ex/Em = 485/535 nm).

### Structural modeling and docking study

An x-ray structure of ABHD10 (AF-Q9NUJ1-F1) or ABAD (PDB: 1U7T) was obtained from the AlphaFold protein structure database or PDB website. The structure files of protein and ligands for docking were prepared using PyMOL 2.3.0 and Chem3D. The docking grid box is generated by the reactive site using AutoDock Vina and exports the grid dimension file. Subsequently, ligands were docked into the binding sites using AutoDock Vina.^78, 79^ Docking procedures were performed using the default setting with 8 exhaustiveness. These obtained ligand-protein complexes were visualized using PyMOL 2.3.0. All these structural modeling figures were generated by PyMOL 2.3.0. And the yellow dashed line represents hydrogen-bonds.

### LC-MS/MS SAMPLE PREPARATION, DATA ACQUISITION, AND ANALYSIS

#### LC-MS/MS sample preparation for tandem mass tag (TMT) chemical proteomics

HEK293T proteomes (427 µL, 2.3 mg/mL) were treated with tested compounds (5 µL, 100×) and incubated for 60 min at RT. Subsequently, 5 µL of either AHL-PuP-2 (100×) or ZH-2-087 (100×) was added to the reaction mixture to yield a final concentration of 100 µM. The mixture was then incubated at RT for 1 h. Following incubation, desthiobiotin-N_3_ (DB-N_3_) was conjugated with probe-modified proteome via a copper-catalyzed azide-alkyne click reaction:^28^ 10 µL DB-N₃ (10 mM), 10 µL TCEP (50 mM), 33 µL TBTA (1.7 mM), 10 µL CuSO₄ (50 mM).

The mixture was incubated at RT for 1 h. The reaction was then quenched by combining it into a MeOH: CHCl_3_: H_2_O mixture (4:1:1.5, 4 mL) into the 2-dram vial to remove excess click reagents. The mixture was centrifuged at 1400 g for 3 min at 4 °C. The top and bottom layers were carefully removed, and the protein layer was transferred to screw-cap microcentrifuge tubes with 600 µL MeOH. The vials were rinsed with 150 µL CHCl3 and 600 µL H2O, respectively. The mixture was centrifuged at 1400 g for 3 min, and the top and bottom layers were discarded. The resulting protein pellet was suspended in 600 µL MeOH and centrifuged at 14000 g for 5 min to remove MeOH.

Following resuspension in 500 µL of 6 M urea/25 mM ammonium bicarbonate (Ambic), 5 µL of Dithiothreitol (DTT, 1M) was added, and the mixture was incubated at 65 °C for 15 min to reduce disulfide bonds. And then 40 µL of Iodoacetamide (IA, 1M) was added, and the reaction was incubated at RT for 30 min in the dark to alkylate the cysteine residues. A second extraction was then performed to remove excess iodoacetamide (IA) and dithiothreitol (DTT), using the same procedure used for the removal of excess click reagents. Proteins were digested overnight with Tryp-Lys-C (7.5 µg, 37 °C) in 25 mM Ambic. The resulting peptides were desalted using Pierce Peptide Desalting Spin Columns in preparation for TMT labeling (ThermoFisher Scientific cat#89852).

### LC-MS/MS sample preparation for tandem mass tag (TMT) chemical proteomics (*in situ*)

HEK293T cells (or corresponding cell lines) were cultured to approximately 90% confluency (∼5 × 10⁶ cells per 100 mm dish) and treated with the tested compounds in serum-free medium for 2 h. Cells were then harvested following the standard protocol described in the “Gel-based ABPP (in situ)” section. HEK293T proteomes (432 µL, 2.3 mg/mL) were incubated with the SuPUR probe AHL-PuP-2 (final concentration of 100 µM). The reaction mixture was incubated at RT for 1 h. Subsequent sample preparation followed the established workflow detailed in the “LC-MS/MS sample preparation for tandem mass tag (TMT) chemical proteomics” section.

### Tandem mass tag (TMT) labeling ^32^

The peptide concentration was determined using Pierce Quantitative Colorimetric Peptide Kit. Peptides were normalized to a concentration of 5 µg/µL in 200 Mm EPPS (pH 8.5). For TMT labeling, 10 µL of peptides were mixed with 5 µL of TMTsixplex isotopomers reagent (100 µg) in PCR tubes, and the reaction was incubated for 1 h at RT with shaking at 500 rpm. The reaction was quenched by adding 1.0 µL of 5% hydroxylamine to each tube, followed by incubation at RT for 15 min with shaking at 500 rpm. Subsequently, 1 µL of formic acid (FA) was added to each tube to acidify and neutralize the excess hydroxylamine. All TMT-labeled samples were combined into one low-binding tube and dried using SpeedVac.

### Enrichment of SuPUR probe modified peptides

Samples were resuspended in 300 µL Optima water, and 5 µL of resuspension was removed for unenriched proteomics analysis (The same LC-MS/MS method used for post-enrichment competitive TMT-ABPP was applied). The resulting samples (295 µL) were incubated with 100 µL of avidin agarose and diluted with DPBS to a final volume of 5.5 mL. The mixture was incubated at RT for 1 h, followed by centrifugation at 1,400 × g for 1 minute to pellet the beads. The supernatant was then carefully decanted. The beads were washed three times with 10 mL of AmBic and an additional three times with 10 mL of Optima-grade water. Probe-modified peptides were eluted by incubating the beads three times with elution buffer (50% acetonitrile, 0.1% formic acid). Eluted samples were snap-frozen and subsequently dried using SpeedVac. Dried samples were resuspended in 50 µL of mobile phase A (0.1% FA in Optima water). Peptide cleanup was performed using StageTips (Octadecyl C18 47 mM, Millipore Sigma, Cat#173427). The peptides were snap-frozen and dried using SpeedVac. Samples were then resuspended in 25 µL of mobile phase A and prepared for LC-MS/MS analysis.

### LC-MS/MS evaluation of TMT-tagged SuPUR-modified peptides

Peptides were analyzed with nano-electrospray ionization-liquid chromatography-mass spectrometry (LC-MS/MS) on a Vanquish Neo UHPLC (ThermoFisher Scientific) coupled to an Orbitrap Eclipse Tribrid mass spectrometer or Vanquish Neo UHPLC (ThermoFisher Scientific) coupled to an Orbitrap Exploris 480 Tribrid mass spectrometer. Peptides were separated an 85 min gradient by reverse phase LC using 3 µm C18 (20 cm) as follows: (A: 0.1% formic acid/H2O; B: 80% ACN, 0.1% formic acid in H2O): 0–2 min 4% B, 450 nL/min; 2–4 min 4.5% B, 450 nL/min; 4–24 min 20% B, 300 nL/min; 24–74 30% B, 300 nL/min; 74–79 min 45% B, 300 nL/min; 79–81 min 50% B, 300 nL/min; 81–83.5 min 99% B, 450 nL/min; 83.5–85 min 99% B, 450 nL/min. Data was collected using a top 30 ddMS2 method using the orbitrap detection for both the MS1 and MS2 spectra. MS1 spectra were acquired at 120K resolution, with automatic gain control set to 125K, and a maximum injection time of 20 ms. The dynamic exclusion was set to 80 seconds (3 times the expected peak width) using an intensity threshold of 8.0E3. MS2 spectra were acquired with the TurboTMT setting enabled, a 15K resolution, quadrupole isolation width of 0.7m/z, with an AGC of 25K, and a max IT of 75 ms. Fragmentation by HCD was achieved at a fixed normalized collision energy of 36%.

### LC-MS/MS data analysis for TMT-tagged SuPUR modified peptides

Peptide identification and quantification was accomplished with Proteome Discoverer 3.0 and a PMI-Byonic node. MS2 spectra were searched against the human protein database (UniProt, download date 01/17/2024) using the following parameters: ≤2 missed cleavages, 10 ppm precursor mass tolerance, 20 ppm fragment mass tolerance, and 1% protein false discovery rate. Modifications considered included SuPUR (+635.2737, Y, K, variable), methionine oxidation (+15.9949, M, variable), cysteine carbamidomethylation (+57.021464, C, fixed), and TMT modification (+229.1629, N-term, K, variable). Peptides used for quantification met the following quality control criteria: PMI-Byonic Score ≥300, delta ppm err. 5, co-isolation interference threshold ≤ 50%, reporter ion S/N ≥10, and were present in n≥2 biological replicates. TMT channels were normalized to the total peptide amount. Volcano plots were generated by grouping PSMs using the peptide isoform node with cutoffs of Log2 FC≥1.0, and p-value ≤0.05. For sites of interest, a median ratio was calculated from all isoforms adducted at the same position.

### LC-MS/MS sample preparation for label-free chemical proteomics

HEK293T proteomes (432 µL, 2.3 mg/mL) were treated with 5 µL of either AHL-PuP-2 (100 ×) or ZH-2-087 (100 ×) to achieve a final concentration of 100 µM. The mixture was incubated at RT for 1 h. Following incubation, desthiobiotin-N_3_ (DB-N_3_) was conjugated to probe-modified proteome via a copper-catalyzed reaction. The subsequent sample preparation followed the standard workflow for “LC-MS/MS sample preparation for tandem mass tag (TMT) chemical proteomics”.

### LC-MS/MS evaluation of label-free SuPUR-modified peptides

Peptides were analyzed with nano-electrospray ionization-liquid chromatography-mass spectrometry (LC-MS/MS) on a Vanquish Neo UHPLC (ThermoFisher Scientific) coupled to an Orbitrap Eclipse Tribrid mass spectrometer or Vanquish Neo UHPLC (ThermoFisher Scientific) coupled to an Orbitrap Exploris 480 Tribrid mass spectrometer. Peptides were separated an 81 min gradient by reverse phase LC using 3 µm C18 (20 cm) as follows: (A: 0.1% formic acid/H_2_O; B: 80% ACN, 0.1% formic acid in H_2_O):0–2 min 4% B, 450 nL/min;2–4 min 4% B, 450 nL/min;4–5 min 20% B, 450 nL/min;5–35 min 30% B, 300 nL/min; 35–60 min 45% B, 300 nL/min;60–65 min 50% B, 300 nL/min; 65–67 min 99% B, 300 nL/min; 67–69 min 99% B, 450 nL/min;69–71 min 4% B, 450 nL/min;71–81 min 4% B, 450 nL/min. Data was collected using a top 30 ddMS2 method using the orbitrap detection for both the MS1 and MS2 spectra. MS1 spectra were acquired at 120K resolution, with automatic gain control set to 125K, and a maximum injection time of 20 ms. The dynamic exclusion was set to 106 seconds (3.3 times the expected peak width) using an intensity threshold of 8.0E3. MS2 spectra were acquired at 15K resolution, quadrupole isolation width of 1.6m/z, with an AGC of 25K, and a max IT of 100 ms. Fragmentation by HCD was achieved at a fixed normalized collision energy of 30%.

### LC-MS/MS data analysis for label-free SuPUR modified peptides

Peptide identification and quantification was accomplished with Proteome Discoverer 3.0 and a PMI-Byonic node. MS2 spectra were searched against the human protein database (UniProt, download date 01/17/2024) using the following parameters: ≤2 missed cleavages, 10 ppm precursor mass tolerance, 20 ppm fragment mass tolerance, and 1% protein false discovery rate. Modifications considered included SuPUR (+635.2737, Y, K, variable), methionine oxidation (+15.9949, M, variable), and cysteine carbamidomethylation (+57.021464, C, fixed). Peptides used site qualification met the following quality control criteria: PMI-Byonic Score ≥300, delta ppm err. 5, were present in n=3 biological replicates.

### Post-enrichment Competitive TMT-ABPP

Samples were prepared as described previously with modifications.^80^ Briefly, fractionated cell lysate (2 mg/mL in 1X DPBS, 500 µL) was treated with FP-biotin (1 mM, 2 h, RT), and excess probe was removed by chloroform:methanol extraction. Protein pellets were resuspended in 500 µL 6M urea/ 25 mM triethylammonium bicarbonate buffer (TEAB), and samples were reduced (10 mM DTT, 15 min, 65°C), alkylated (40 mM IAA, 30 min, RT), and then diluted with 1X DPBS and SDS (2% final) for a total volume of 500 µL. Biotinylated proteins were enriched with avidin agarose beads (100 µL) and incubated for 1 h at RT while rotating. The beads were washed with 1X DPBS and LC-MS grade water, then transferred to a low-bind microcentrifuge tube and pelleted to remove excess water. The beads were resuspended in 200 µL of EPPS buffer (200 mM, pH 8.5) and on-bead digestions were performed for 12 h at 37 °C with sequence-grade modified trypsin (Promega; 2 µg total). Tryptic peptides were collected by washing avidin beads with 150 µL EPPS buffer over a Bio-Rad biospin column. Samples were dried by SpeedVac, desalted by C18-StageTip, and resuspended in 10µL of EPPS (200 mM, pH 8.5). Each sample was labeled with 100µg TMTsixplex^TM^ for 1h at RT with shaking (500 rpm). The reactions were quenched with 1 µL 5% hydroxylamine for 15min. All 6 channels were combined, dried, and resuspended in 50 µL 0.1% formic acid. Samples were cleaned by StageTip, dried, and resuspended in 50 µL 0.1% formic acid and stored at −80°C until LC-MS analysis.

### Post-enrichment Competitive TMT-ABPP (*in situ*)

HEK293T cells were cultured to approximately 90% confluency (∼5 × 10⁶ cells per 100 mm dish) and treated with the tested compounds in serum-free medium for 2 h. Cells were then harvested following the standard protocol described in the “Gel-based ABPP (*in situ*)” section. HEK293T proteomes (500 µL, 2.0 mg/mL) were incubated with the FP-biotin probe (5 µL, 1 mM). The reaction mixture was incubated at RT for 2 h. Subsequent sample preparation followed the established workflow detailed in the “Post-enrichment Competitive TMT-ABPP” section.

### Data acquisition

LC-MS/MS evaluation of TMT-modified peptides was performed with a Vanquish Neo UHPLC coupled to an Orbitrap Exploris 480 mass spectrometer. Peptides were separated on an 140 min gradient by reverse phase LC using 3 µm C18 (20 cm) as follows: (A: 0.1% formic acid in H_2_O; B: 80% ACN, 0.1% formic acid in H_2_O: 0–2 min 4% B, 450 nL/min; 2–2.5 min 5% B, 300 nL/min; 2.5–78 min 25% B, 300 nL/min; 78–123 45% B, 300 nL/min; 123–124.973 min 99% B, 450 nL/min; 124.973-127.604 min 99% B, 450 nL/min; 127.604-129.577 min 1% B, 450 nL/min; 129.577-140 min 1% B, 450 nL/min). Data was acquired using a top 30 ddMS2 method. MS1 spectra were acquired at 120K resolution with a max IT of 20 ms and MS2 spectra were taken with an AGC of 25K, quadrupole isolation width of 0.7 m/z, and max IT of 50 ms.

### Data analysis

Peptide identification and quantification for TMT-tagged peptides was performed using Proteome Discoverer 3.0 housing a PMI-Byonic node. MS2 spectra were searched against the human protein database (UniProt, download date 01/17/2024) using the following parameters: ≤ 2 missed cleavages, 10 ppm precursor mass tolerance, 20 ppm fragment mass tolerance, and 1% protein false discovery rate. Modifications considered included methionine oxidation (+15.9949, M, variable), cysteine carbamidomethylation (+57.021464, C, fixed), and TMT modification (+229.1629, N-term, K, variable). Peptides used for protein quantification met the following quality control criteria: PMI-Byonic Score ≥300, delta ppm err. 5, co-isolation interference threshold ≤ 50%, reporter ion S/N ≥10, TMT-modified, and were present in n = 2 biological replicates. Proteins were quantified had 3 or more unique, quality peptides matched. TMT channels were normalized to total peptide amount. Volcano plots were generated based on protein abundance ratios (inhibitor/vehicle) with cutoffs of Log_2_ FC ≥ 0.5, and p-value ≤ 0.05.

### Predicting druggable pockets

Druggable sites for proteins of interest were mined using the predictions within canSAR.ai, and that underlying methodology has been detailed elsewhere.^81^ Summarily, all structures in the Protein Databank were submitted to pocket finding using SURFNET,^82^ and 26 geometric and thermodynamic features were computed for each pocket to provide descriptors of each pocket. “Drug-like” ligands within the PDB were taken as those with physicochemical parameters consistent with expanded Lipinski Rule of 5^83^ criteria, additionally considering topological polar surface area (≤ 140 Å) and the number of rotatable bonds (≤ 10).^84^ Pockets containing these ligands were labeled as “druggable” positives, and all other pockets were left as unlabeled. A two-step iterative Positive Unlabeled Learning scheme employing the spy technique^85^ was employed using a Random Forest to assign labels to reliable negatives. A final Random Forest classifier was trained on the dataset of positives and reliable negatives and deployed to canSAR.ai, with predictions made on new accessions weekly.

### Gene ontology (GO) analysis

Gene Ontology (GO) enrichment analysis was performed, and the results were visualized with the OmicShare tool, an online platform for data analysis. https://www.omicshare.com/tools/Home/Soft/enrich_circle.^60^

Gene Ontology (GO) is an internationally standardized system for the functional classification of genes. It offers dynamically updated, controlled vocabulary for comprehensively describing the attributes of genes and gene products across species. In GO analysis, each functional attribute is represented by a GO term, which serves as a basic unit within the ontology. GO functional analysis includes both the annotation of differentially expressed genes (DEGs) with GO terms and the enrichment analysis of these terms.

DEGs are annotated with Gene Ontology (GO) terms using the GO database (http://www.geneontology.org/), and the number of genes associated with each term is quantified. Subsequently, a hypergeometric test is performed to identify GO terms that are significantly enriched in the DEG set relative to the genomic background.

The p-value for each GO term is calculated using the following formula:

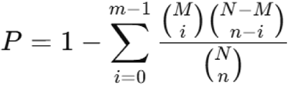

Where:

N is the total number of genes with GO annotations (background set),

*n* is the number of DEGs within N,

M is the number of genes annotated with a specific GO term in N,

*m* is the number of DEGs annotated with that specific GO term.

The resulting p-values are corrected for multiple testing using the False Discovery Rate (FDR) method. GO terms with a corrected p-value ≤ 0.05 are considered significantly enriched among the DEGs.

### Kyoto encyclopedia of genes and genomes (KEGG) analysis

Kyoto Encyclopedia of Genes and Genomes (KEGG) enrichment analysis was performed and results were visualized using the OmicShare tool (https://www.omicshare.com/tools/home/report/koenrich.html). ^63^

The KEGG is a widely used public database that offers comprehensive information on biological pathways. To identify pathways significantly enriched with DEGs, this enrichment analysis was performed based on KEGG database. The hypergeometric test was employed to determine whether a given pathway was statistically overrepresented among the DEGs, compared to the genomic background.

The P-value was calculated using the following formula:

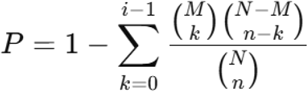

N is the total number of background genes,

*n* is the number of DEGs,

M is the number of background genes annotated to a specific pathway,

*i* is the number of DEGs annotated to the pathway.

The P-values were adjusted for multiple testing using the FDR method. Pathways with a corrected P-value ≤ 0.05 were considered significantly enriched among the DEGs.

### Disease ontology (DO) analysis

Disease ontology (DO) enrichment analysis was performed and results were visualized using the OmicShare tool (https://www.omicshare.com/tools/Home/Soft/doenrichsenior).^60^

DO database is a curated resource designed to describe gene functions related to human diseases. The objective is to provide the biomedical community with a consistent and standardized vocabulary for human disease terms, phenotypic features, and associated medical concepts. Similar to GO, DO integrates multiple disease classification systems such as MeSH and ICD, providing a unified framework for both common and rare human diseases. To identify disease terms significantly enriched among DEGs, enrichment analysis was performed based on Disease Ontology. The hypergeometric test was applied to assess whether a specific Disease Ontology ID (DOID) was statistically overrepresented in the list of DEGs compared to the genomic background.

The P-value was calculated using the following formula:

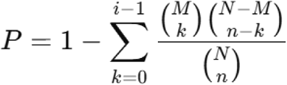

N is the total number of background genes,

*n* is the number of DEGs,

M is the number of background genes annotated to a specific DOID,

*i* is the number of differentially expressed genes annotated to that DOID.

P-values were adjusted for multiple testing using the FDR method. DO terms with a corrected P-value ≤ 0.05 were considered significantly enriched among the DEGs.

### Domain enrichment analysis

Probe-modified sites were matched to annotated PROSITE domains in humans (available on EXPASY, https://ftp.expasy.org/databases/prosite/). All probe-modified sites falling with the range of each annotated domain were considered a single ‘hit’ for AHL-PuP-2 or ZH-2-087. The natural frequency of each domain in humans, *P(D)*, was calculated by summing each instance of a domain, *n(D)*, and dividing by the total number of annotated domains, *N*:

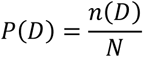

Frequencies of probe-modified domains, *P(D)’*, were calculated similarly by summing each hit within a domain class, *n(D)’*, and dividing by the total number of matched domains, *N’*:

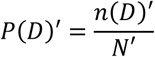

Fold Enrichment values were calculated by dividing the probe-modified domain frequency over its frequency in nature:

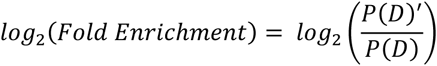

P-values for matched domains were calculated using a two-sided binomial test (binomtest function in SciPy) where *n* is the number of matched domains, and *k* is how many times a specific domain appears in the matched dataset:

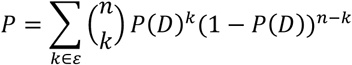

The P-values were corrected by applying a Benjamini-Hochberg correction and filtered using a 5% false discovery rate (FDR). ProSite domains with fold enrichment > 1, Q-Value < 0.05) were considered significantly enriched.

### DrugBank and ChEMBL analysis

UniProt accession IDs identified by AHL-PuP-2 were submitted to the UniProt Retrieve/ID Mapping tool to retrieve corresponding DrugBank and ChEMBL identifiers for subsequent overlap analysis. https://www.uniprot.org/id-mapping.

### Purinome analysis

Purinome protein targets were identified using UniProt database, focusing on reviewed entries annotated with purine-based cofactors and substrates, including ATP/GTP, NAD/NADP, CoA, SAM, FAD/FADH2, and GMP/AMP. https://www.uniprot.org/uniprotkb.

### Statistical analysis and reproducibility

Data are presented as the mean ± s.d. from independent experiments. All statistical analyses were performed on n = 3 biologically independent experiments, unless otherwise noted in the figure captions, using GraphPad Prism 10.0 software.

### Data availability

All crystal structures for small molecules have been deposited in the CCDC database (https://www.ccdc.cam.ac.uk/profile/). The deposition numbers are listed in Supplementary **Table 6**. Co-crystal and apo structures of ACAT2 have been verified and deposited in the Protein Data Bank (PDB) with the accession code 9V13 and 9V37. Structural model files have been deposited at Figshare (https://figshare.com/s/ec20e5585019897814f2). Proteomics data was deposited at ProteomeXchange via the PRIDE database (PXD: 066797) and are publicly available as of the date of publication. Reviewer can access the dataset by logging in to the PRIDE website using the following account details: Username. Password: CqnItidUGBY9. This paper does not report the original code.

## Acknowledgements

We thank all members of the Hsu lab for helpful discussions and review of the manuscript. This work was supported by a Recruitment of Rising Stars Award from CPRIT (RR220063 to K.-L.H.), the Mark Foundation for Cancer Research (Emerging Leader Award to K.-L.H.), and a Research Grant Award from The Welch Foundation (F-2143-20230405 to K.-L.H.). We thank the X-ray Diffraction Laboratory at the University of Texas at Austin for their assistance with the crystallization of SuPUR ligands. We thank the experimental facility and the technical services provided by the National Synchrotron Radiation Research Center (NSRRC), a national user facility supported by NSTC, Taiwan.

## Author contributions

K.-L. H. and Z. L. conceived and directed the project and wrote the manuscript. K.-L. H. supervised this project. All authors edited the revised manuscript. Z. L. performed most of the experiments. W.W. performed domain enrichment analysis. C. C. and H. T. performed the protein crystal structure experiments. A.-H. L. helped with SuPUR chemistry early in the project. M.-W. F., O.-L. M., M.-L. W., and D.-M. L. developed LC-MS/MS methods. P. G., and B. L provide the druggable sites database.

## Competing interests

K.-L.H. is a founder and scientific advisory board member of Hyku Biosciences. A patent application has been filed on the work presented in this manuscript.

## Additional information

The online version contains supplementary material available at https://doi.org/xxx. Correspondence and requests for materials should be addressed to Ku-Lung Hsu.

### Supplementary information

**Supplementary Table 1** Label-free proteomics data

**Supplementary Table 2** Data enrichment analysis

**Supplementary Table 3** Competitive TMT-ABPP proteomic data

**Supplementary Table 4** Non-enriched proteomics data

**Supplementary Table 5** Data collection, phasing, and refinement statistics for ACAT^apo^ and ACAT2^ZH-2-077^ structures

**Supplementary Table 6** Key resources

**Supplementary Figs. 1-4** and compounds characterization data

**Source data**

Unprocessed gel images.

### Extended data

**Extended data Fig. 1.**
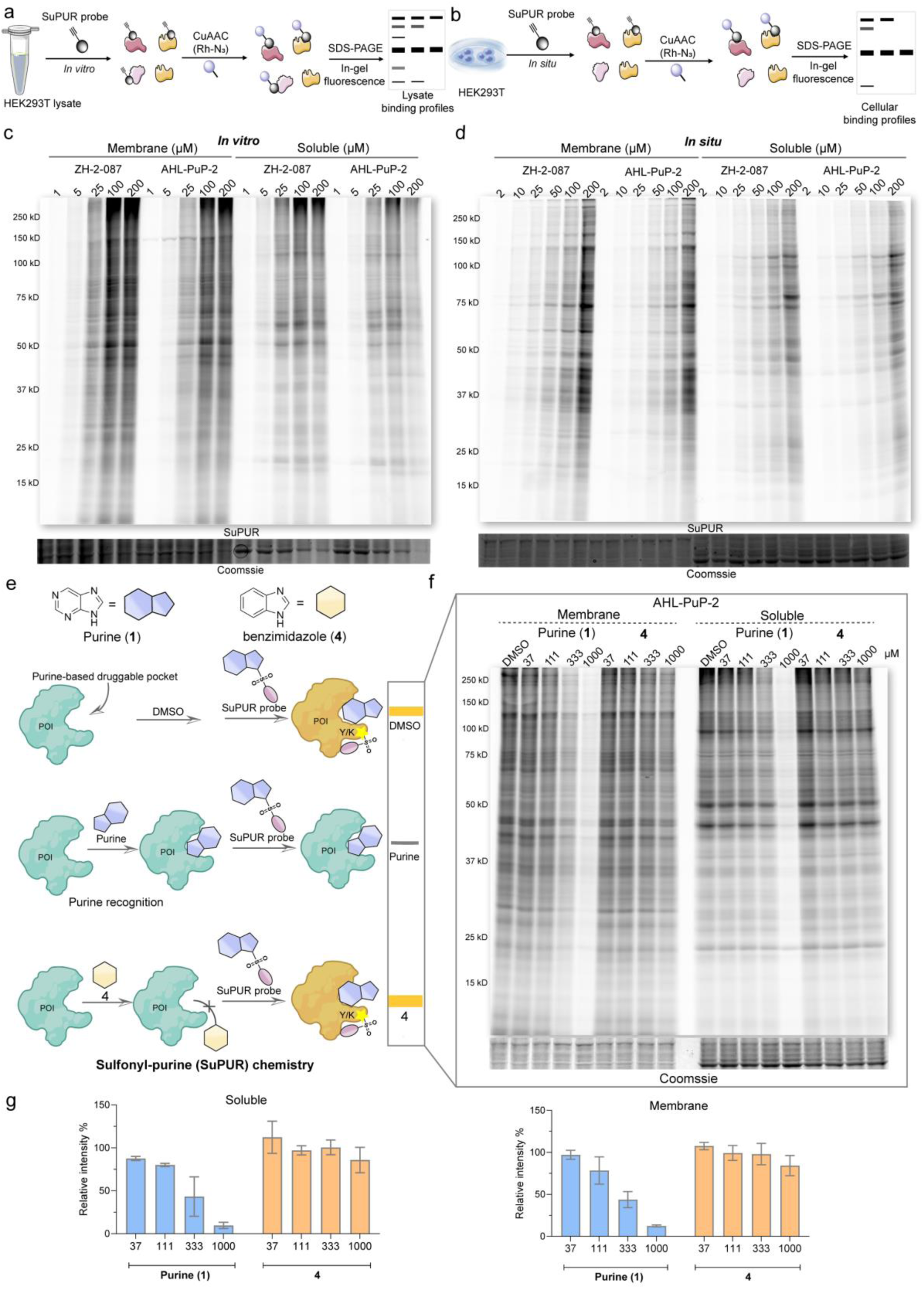
Developing SuPUR chemistry to target the human purine interactome. **a** and **b,** Gel-based activity-based protein profiling (ABPP) workflow by *in vitro* labeling (**a**) and *in situ* labeling (**b**). (n = 3). **c,** Proteins labeled by ZH-2-087 and AHL-PuP-2 in lysates were separated by SDS-PAGE (*in vitro*, 1 h). Reduced band intensity at the highest concentration (200 µM) is due to the probes crashing out of solution. (n = 3). **d,** Proteins labeled by ZH-2-087 and AHL-PuP-2 in cells were separated by SDS-PAGE (*in situ*, 2 h). (n = 3). **e,** The proposed SuPUR reaction mechanism involves covalent targeting of the protein of interest (POI), with purine serving as a crucial recognition group for covalent binding. **f,** Competitive gel-based ABPP analysis of purine (**1**) and control benzimidazole (**4**) in HEK293T proteomes using FP-Rh (*in vitro*, rt, 1 h). **g,** Quantification of gel bands was performed using Bio-Rad Image Lab 6.0. The results showed that purine exhibited significant competition of SuPUR probe labeling, whereas the benzimidazole control, showed no detectable inhibitory activity. Data are presented as mean ± SD (n = 3).

**Extended data Fig. 2.**
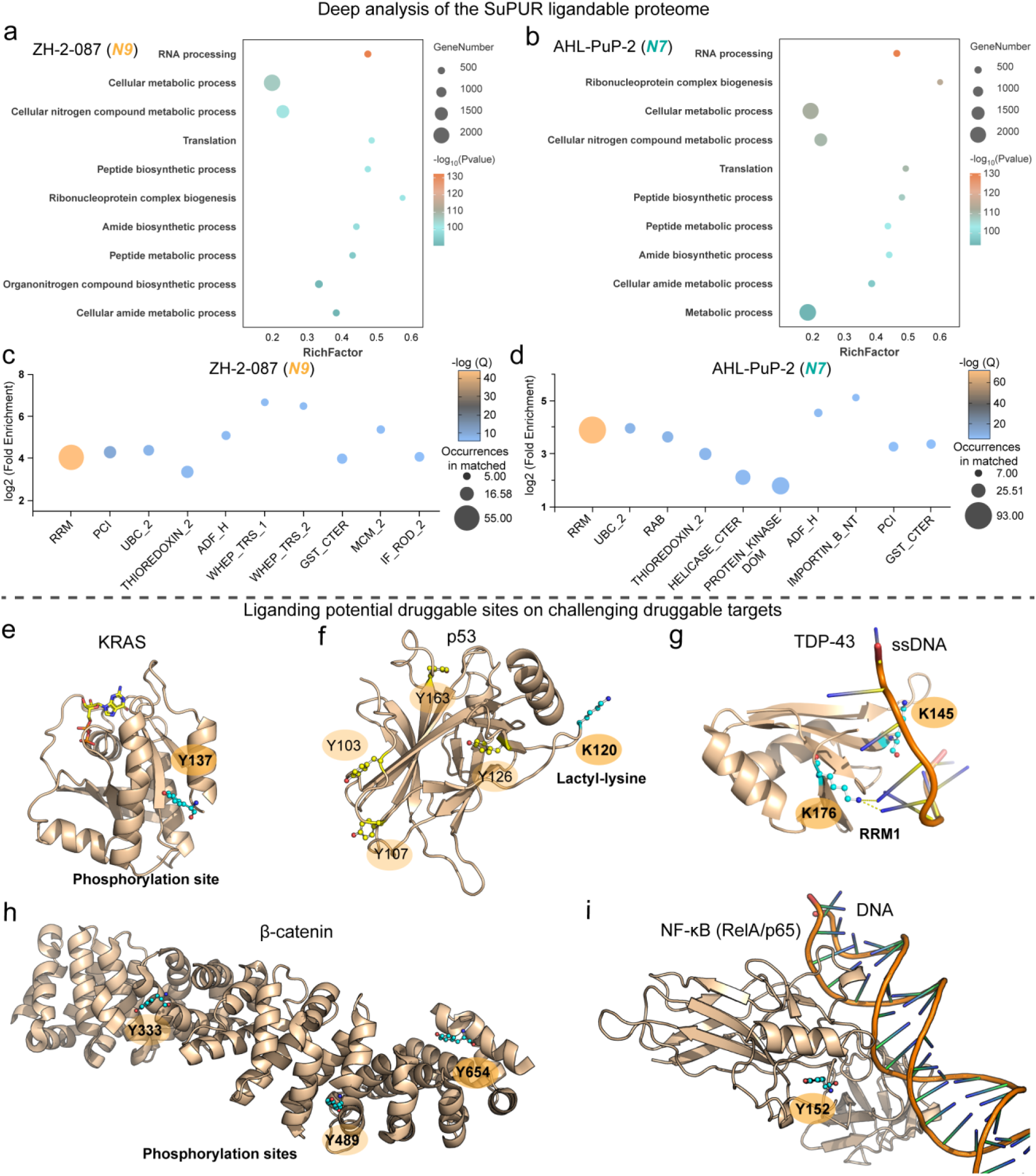
SuPUR chemoproteomics establishes the targetable human purine interactome. **a** and **b,** Gene ontology (GO) enrichment analysis of ZH-2-087 (**a**) and AHL-PuP-2 (**b**) modified proteins in the HEK293T proteome for biological process; the top 10 enriched GO terms are shown. **c** and **d,** Domain enrichment analysis of SuPUR sites identified by ZH-2-087 (**c**) and AHL-PuP-2 (**d**) in HEK293T proteome. **e,** Representative sites targeted by SuPUR chemoproteomics on intractable proteins of interest (POI). KRAS Tyr137 (Y137) is a conserved residue and a potential phosphorylation site (PDB: 5VQ2).^86^ Targeting Y137 through site-specific liganding may offer a strategy to modulate KRAS enzymatic activity.^87^ **f,** p53 Lys120 (K120) is in the DNA-binding domain (DBD) that undergoes acetylation (PDB: 1UOL),^88^ regulating both transcription-dependent and transcription-independent apoptotic pathways.^89^ **g,** TDP-43 is primarily an RNA-binding protein, with Lys145 (K145) and Lys176 (K176) located in the RNA Recognition Motif 1 (RRM1) domain (PDB: 4IUF).^88^ Notably, K145 forms a hydrogen bond interaction with single-stranded DNA (ssDNA). These lysine residues represent potential regulatory hotspots; therefore, targeting K145 and K176 through site-specific ligand engagement could provide a strategy to modulate TDP-43 activity.^88^ **h,** SuPUR modified sites are known phosphorylation sites on β-catenin (PDB: 1JDH).^90^ Phosphorylation at tyrosine 654 has been shown to enhance Wnt signaling and promote intestinal tumorigenesis.^91^ **i,** Tyr152 (Y152) on NF-κB (RelA/p65) is located within the DNA-binding interface (PDB:2O61),^92^ suggesting a potential regulatory role in modulating DNA interaction and transcriptional activity.

**Extended data Fig. 3.**
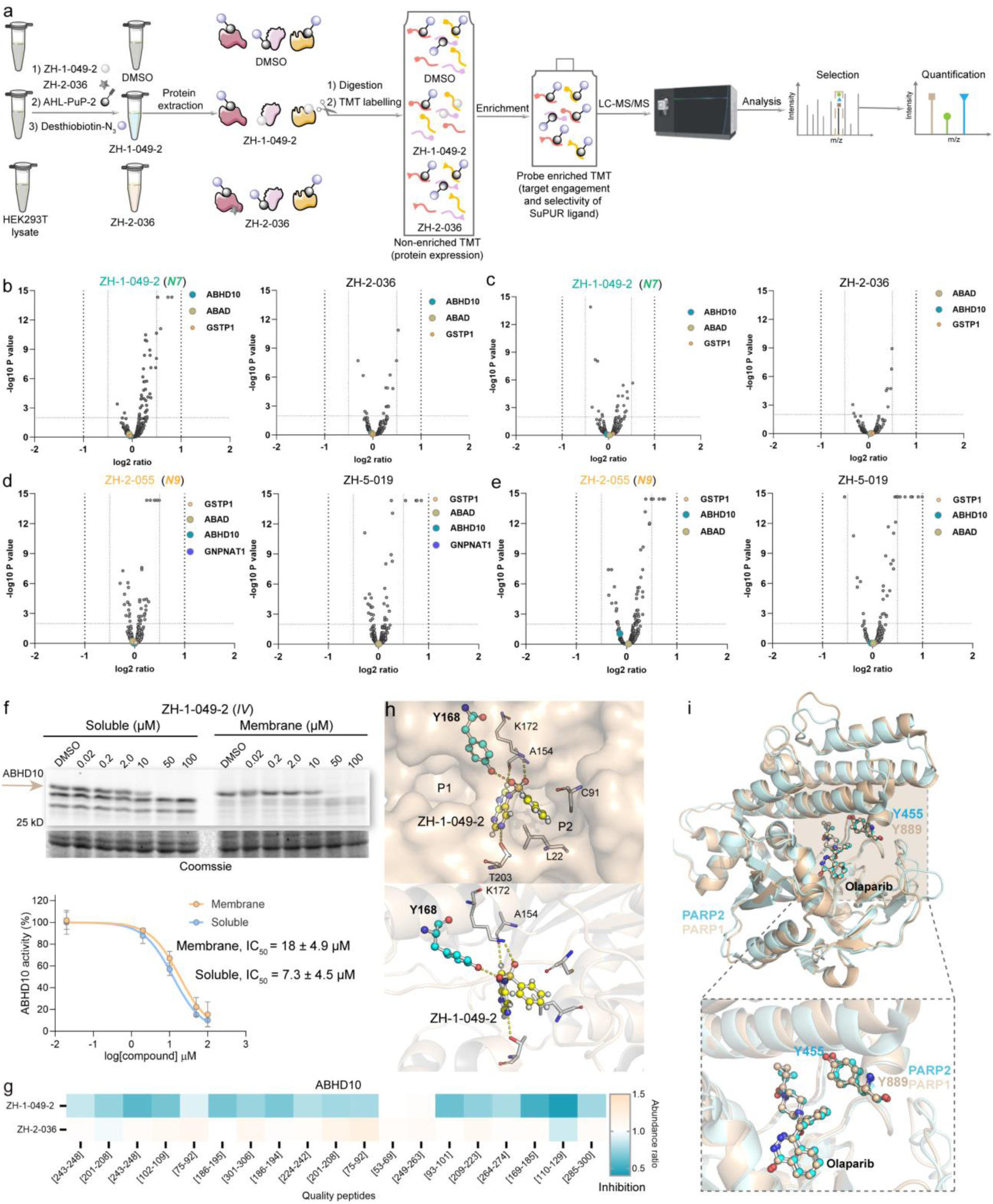
Lead SuPUR ligands demonstrate site-specific and regio-selective protein modulation. **a,** Workflow of competitive SuPUR TMT-ABPP where the SuPUR probe-modified peptides are enriched by avidin beads and blockade by SuPUR ligands is indicative of target and site engagement. Inclusion of N- delete controls help discern specific from non-specific binding events. **b** and **c,** TMT non-enriched proteomics to evaluate protein expression changes from ZH-1-049-2 (25 µM) or ZH-2-036 (25 µM) treatments in HEK293T proteomes (*in vitro*, rt, 1 h). Volcano plots show the distribution of global soluble proteome (**b**) and membrane proteome (**c**) in comparison between the compound- and the vehicle-treated groups. The results indicated that the protein levels of ABHD10, ABAD, and GSTP1 were not changed compared to the vehicle group. Additionally, GNPNAT1 and PARP2 were not detectable by non-enriched proteomics. The proteins represent the combined results from n = 3 independent biological replicates. **d** and **e,** TMT non-enriched proteomics to evaluate protein expression changes from ZH-2-055 (25 µM) and ZH-5-019 (25 µM) treatments in HEK293T proteomes (*in vitro*, rt, 1 h). Volcano plots show the distribution of global soluble proteome (**d**) and membrane proteome (**e**) in comparison between the compound-treated and the vehicle-treated groups. The results indicated that the protein levels of ABHD10, ABAD, GNPNAT1, and GSTP1 were consistent with those observed in the vehicle group. PARP2 was not detectable in the non-enriched proteomics dataset. The proteins represent the combined results from n = 3 independent biological replicates. **f,** Competitive gel-based ABPP analysis of ZH-1-049-2 in HEK293T proteomes using FP-Rh (*in vitro*, rt, 1 h). Data shown are mean ± SD (n = 3). **g,** Quantified peptides corresponding to ABHD10 were detected in the ZH-1-049-2 and ZH-2-036 treatment groups (*in vitro*, rt, 1 h), with values normalized to vehicle group. (n = 2). **h,** Docking of ZH-1-049-2 into ABAD protein structure (PDB: 1U7T) revealed the ligand occupies the NAD+ pocket and forms hydrogen-bonds with K172 and Y168. Docking was performed with AutoDock Vina.^43^ The figure was generated using PyMOL 2.3.0. **i,** Structural superimposition of catalytic domains of PARP2 (PDB: 8HLJ, pale green) and PARP1 (PDB: 7AAD, wheat) reveals a high degree of similarity in their substrate-binding pockets,^44, 45^ with a root-mean-square deviation (RMSD) of 1.0. The SuPUR liganded site on PARP2 Y455 is conserved in PARP1 but the analogous site (Y889) on PARP1 is not modified by SuPUR ligands.

**Extended data Fig. 4.**
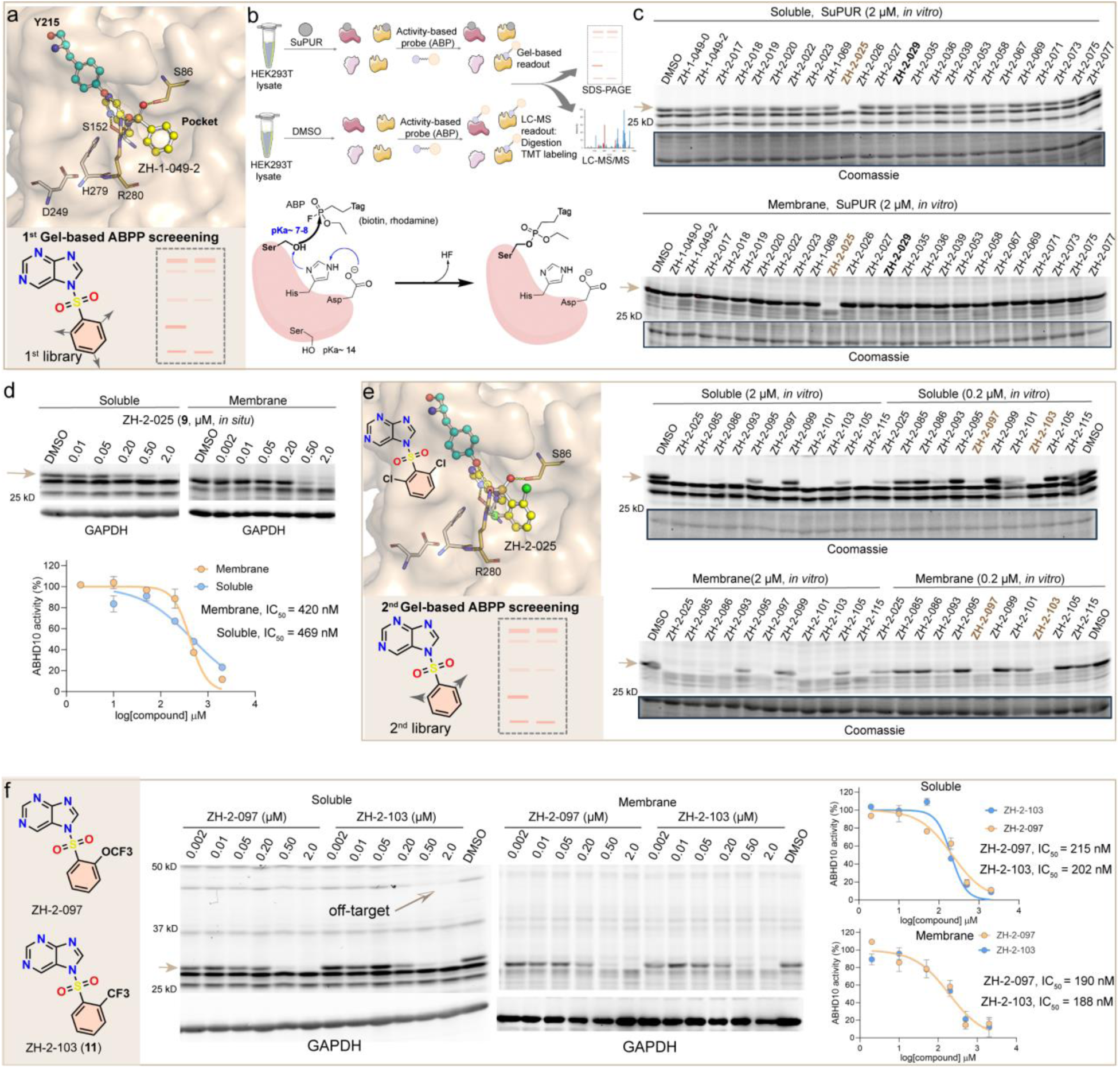
Fragment-based ligand discovery of SuPUR ligands of ABHD10. **a,** Docking of ZH-1-049-2 into human ABHD10 (AF-Q9NUJ1-F1) supports ZH-1-049-2 binding in the serine catalytic pocket and hydrogen-bond interactions (yellow dashes) with Y215, S86, S152, and R280. Docking was performed with AutoDock Vina. The figure was generated using PyMOL 2.3.0. The rational design of the 1^st^ SuPUR library is based on the binding model of ZH-1-049-2 with ABHD10, with an optimization strategy focused on improving occupancy of the catalytic pocket through modification to the terminal phenyl group. **b,** Basic ABPP workflow using fluorophosphonate (FP) probes. The mechanism of action involves covalent modification of the conserved catalytic serine residue in SHs by the FP probe. **c,** 1^st^ FP-Rh-mediated competitive gel-based ABPP screening (Supplementary Fig. 2) was performed in HEK293T proteomes (1 h, rt, *in vitro*), leading to discovery that ZH-2-025 (**9**) exhibited potent inhibitory activity against ABHD10. **d,** Potency of ZH-2-025 against ABHD10 using FP-Rh-mediated competitive gel-based ABPP in HEK293T soluble proteome (IC_50_ of ∼400-500 nM). Data shown are mean ± SD (n = 2). **e,** Docking of ZH-2-025 into human ABHD10 (AF-Q9NUJ1-F1) supports the chloride on the terminal phenyl group could form a hydrogen-bond with R280, potentially enhancing the binding affinity between ZH-2-025 and ABHD10. The rational design of the 2^nd^ SuPUR library (Supplementary Fig. 2) involves the introduction of minor substitutions (electron-withdrawing/donating groups) on the terminal phenyl group, aiming to improve the binding affinity of SuPUR compounds with ABHD10. 2^nd^ FP-Rh-mediated competitive gel-based ABPP screening was performed, indicating that compounds with electron-withdrawing group substitutions enhanced the binding affinity to ABHD10, particularly for ZH-2-097 and ZH-2-103. **f,** The binding activities of ZH-2-097 and ZH-2-103 (**11**) were assessed using FP-Rh-mediated competitive gel-based ABPP in HEK293T proteomes (*in situ*, 2 h). ZH-2-103 shows notable off-target activity against a 50 kDa SH protein. Data shown are mean ± SD (n = 2).

**Extended data Fig. 5.**
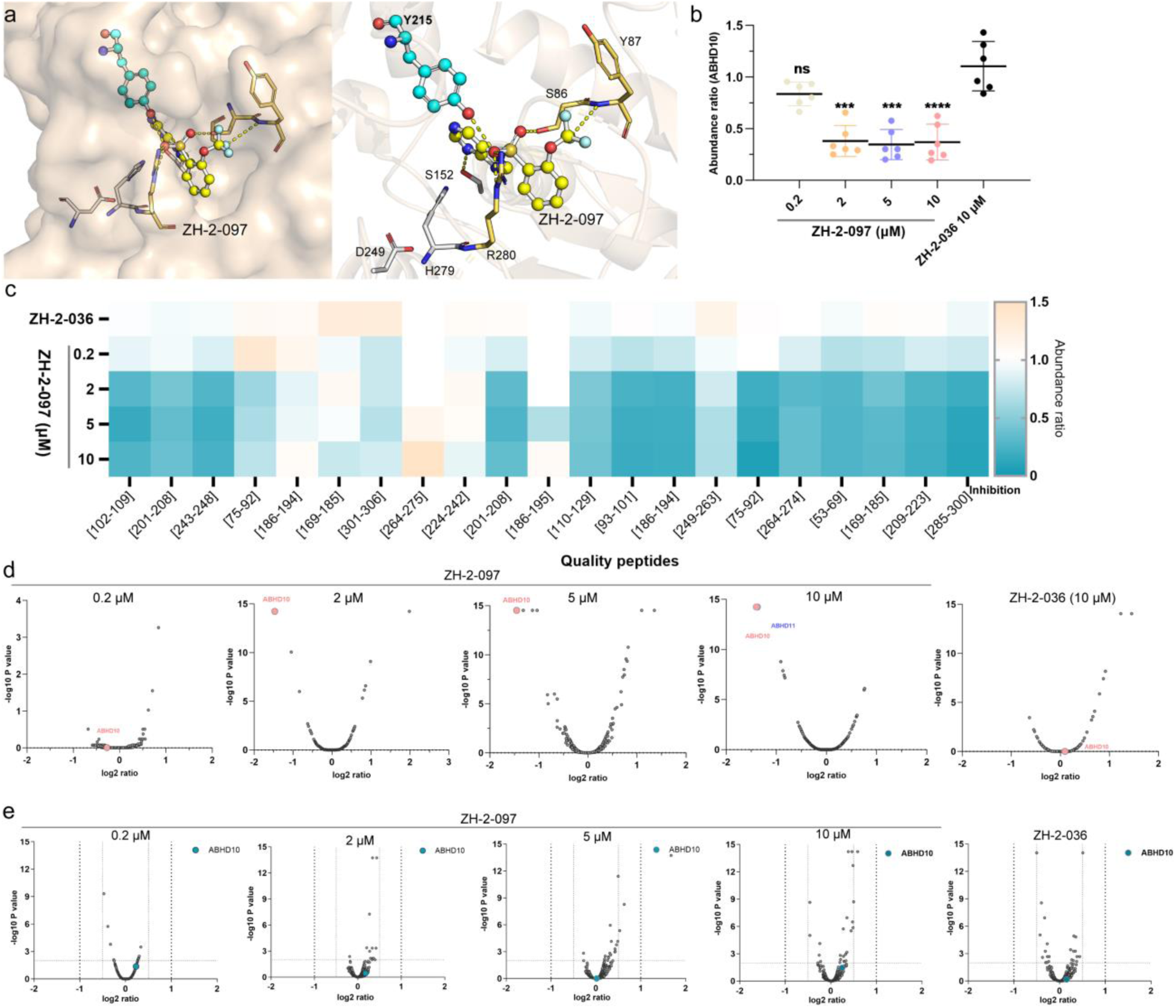
Disrupting the catalytic site of ABHD10 via ZH-2-097 liganding of Y215. **a,** Docking of ZH-2-097 into human ABHD10 (AF-Q9NUJ1-F1) supports ZH-2-097 binding in the serine catalytic pocket and hydrogen-bond interactions (yellow dashes) with Y215, Y87, S86, S152, and R280. Docking was performed with AutoDock Vina. The figure was generated using PyMOL 2.3.0. **b,** The levels of FP-biotin enriched ABHD10 protein were normalized to the DMSO vehicle control group (*in situ*, 2 h), p values were analyzed by one-way ANOVA comparing with the control group ZH-2-036, ***p < 0.001, ****p < 0.0001. Data shown are mean ± SD. (n = 3) **c,** Quantified peptides corresponding to ABHD10 were detected in the ZH-2-097 and ZH-2-036 treatment groups (*in situ*, 2 h), with values normalized to DMSO vehicle group. (n = 3). **d,** FP-biotin-mediated competitive TMT-ABPP analysis of HEK293T cells treated with ZH-2-097 (0.2-10 µM) and ZH-2-036 (N-delete control) versus DMSO vehicle revealed selective inhibition of ABHD10 across all proteins detected in the dataset. The FP-biotin-enriched proteins represent the combined results from n = 3 independent biological replicates **e,** TMT non-enriched proteomics of ZH-2-097 and ZH-2-036 with AHL-PuP-2. Volcano plots show the distribution of global protein expression comparing between the compound-treated and DMSO vehicle groups in HEK293T soluble proteome (*in situ*, 2 h). ABHD10 is highlighted in blue. Additionally, ABHD10 was not detected in the non-enriched proteomics analysis of HEK293T membrane proteome. The proteins represent the combined results from n = 3 independent biological replicates.

**Extended data Fig. 6.**
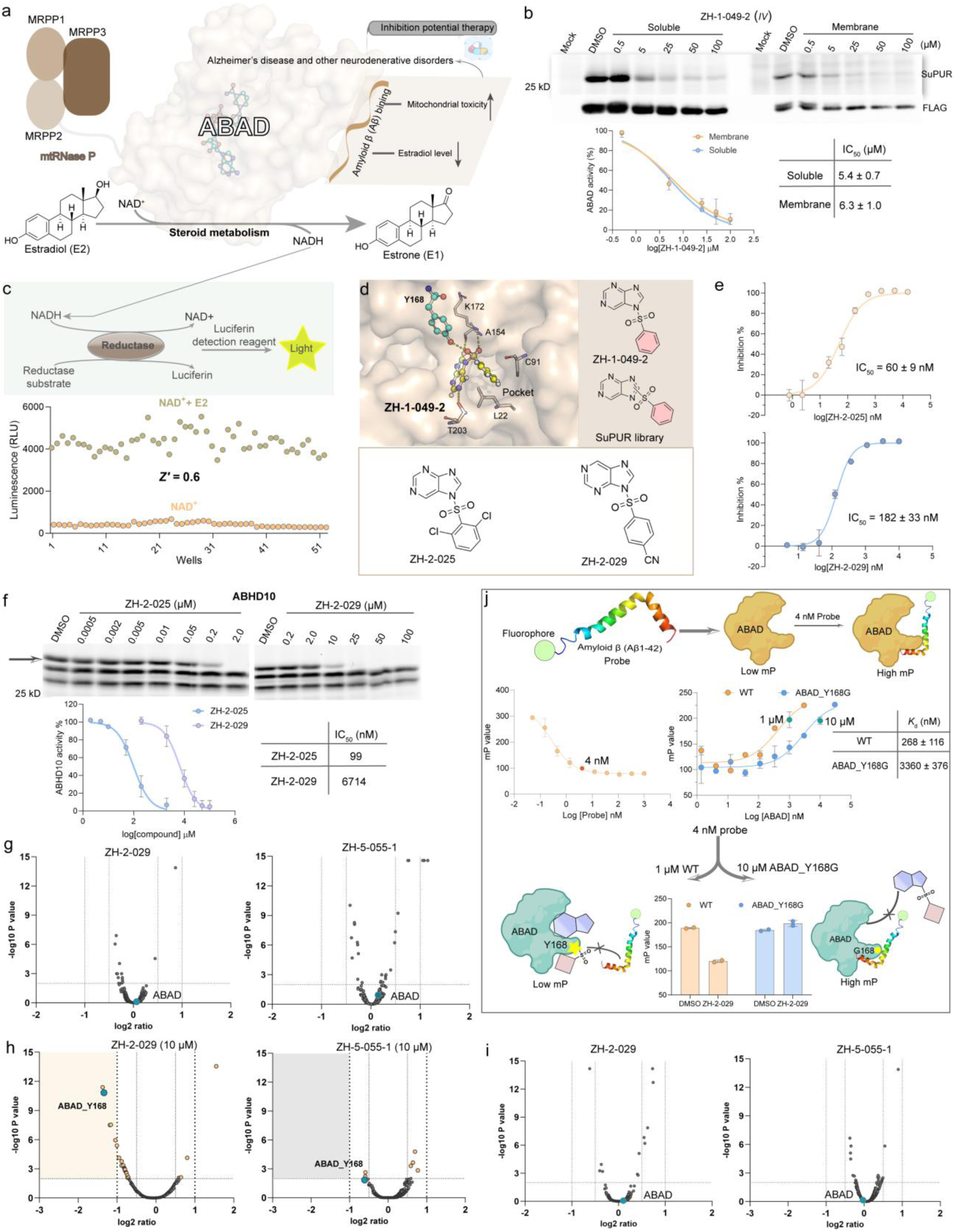
Disrupting catalytic and PPI activity of ABAD using a Y168-specific and N9-regioselective SuPUR ligand. **a,** Aβ-binding alcohol dehydrogenase (ABAD), also known as MRPP2, is a key enzyme involved in steroid metabolism with NAD+ as a cofactor. Aβ binds intracellularly to ABAD, inhibiting its enzymatic activity. The loss of enzymatic function triggers a natural compensatory mechanism, resulting in increased expression of the ABAD gene and elevated levels of the ABAD protein. Subsequently, over-expressed ABAD causes mitochondrial toxicity and reduces estradiol. These are closely related to Alzheimer’s disease and cancers, and inhibiting ABAD could present a broad therapeutic potential for Alzheimer’s disease and cancer.^51^ **b,** The binding activity of ZH-1-049-2 was evaluated using AHL-PuP-2-mediated competitive gel-based ABPP (*in vitro*, 1 h) in HEK293T proteome. Data shown are mean ± SD (n = 3). **c,** Establishment of a NAD(P)/NAD(P)H-Glo assay for ABAD inhibitor screening, along with a schematic representation of the assay mechanism. To evaluate the robustness of the NAD(P)/NAD(P)H-Glo assay, the Z’ factor was determined based on multiple biological replicates. A Z’ factor of 0.6 indicates good assay quality, making this ABAD biochemical assay suitable for high-throughput screening. **d,** The binding model of ZH-1-049-2 with ABAD provides insights into the design of the SuPUR ligand library aimed at fully occupying the NAD+ pocket to enhance binding affinity. Screening of a SuPUR ligand library using the NAD(P)/NAD(P)H-Glo assay identified ZH-2-029 and ZH-2-025 as potent ABAD inhibitors. **e,** Potency of ZH-2-025 and ZH-2-029 against ABAD were determined by NAD(P)/NAD(P)H-Glo assay. Data shown are mean ± SD (n = 3). **f,** The off-target binding activity of ZH-2-025 and ZH-2-029 against ABHD10 was determined using FP-Rh-mediated competitive gel-based ABPP in HEK293T soluble proteome (1 h, *in vitro*). Data shown are mean ± SD (n = 2). **g,** TMT non-enriched proteomics show global protein expression was largely unaffected in HEK293T cells treated with ZH-2-029 or ZH-5-055-1 (N-delete control; 2 h, *in situ*) at 10 µM. ABAD is highlighted in blue color. The depicted proteins represent the combined results from n = 3 independent biological replicates. **h,** Competitive SuPUR TMT-ABPP shows ABAD Y168 is the most significantly liganded site across thousands of quantified AHL-PuP-2 modified peptides in HEK293T membrane proteomes from treated cells (2 h, *in situ*). The AHL-PuP-2 modified peptides represent the combined results from n = 3 independent biological replicates. **i,** TMT non-enriched proteomics shows ABAD and other proteins detected did not show expression changes with ZH-2-029 or ZH-5-055-1 treatments in membrane proteomes from treated cells (2 h, *in situ*) at 10 µM. ABAD is highlighted in blue color. The proteins shown represent the combined results from n = 3 independent biological replicates. **j,** Development of a fluorescence polarization (FP) assay to measure ABAD binding to a Fluor 488-labeled Aβ-peptide (probe). Optimization identified a probe concentration of 4 nM to be suitable for the assay. The enzyme concentrations for ABAD wild-type and Y168G mutant were identified as 1 and 10 µM, respectively, to be optimal for the assay. A significant increase in millipolarization (mP) signal was observed upon incubation of the probe with ABAD and used to determine the binding affinity of Aβ-peptide with ABAD wild-type compared with Y168G mutant. The Y168G mutant showed a ∼10-fold reduction in binding affinity with the Aβ-peptide. Pretreatment with ZH-2-029 resulted inhibition of ABAD-Aβ-peptide interactions as measured by a decreased mP signal. This sensitivity to ZH-2-029 inhibition was lost in the ABAD Y168G mutant. Data shown are mean ± SD (n = 3).

**Extended data Fig. 7.**
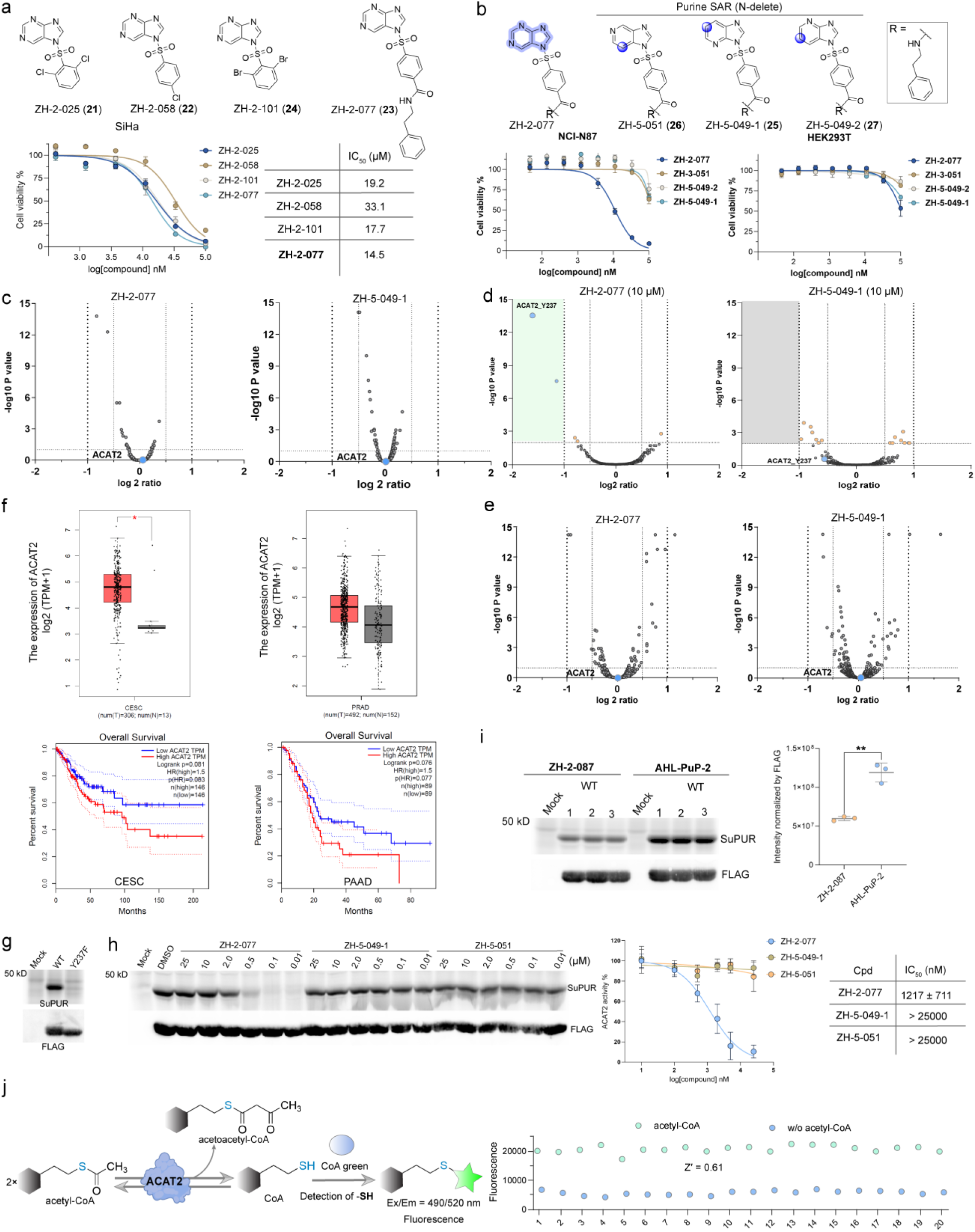
Targeted anti-cancer activity using a Y237-specific and N7-regioselective SuPUR inhibitor of ACAT2. **a,** Determining potency (IC_50_) of hit compounds (>60% inhibition at 30 µM in phenotypic screen) using a luminescent cell viability assay in SiHa cells. Data shown are mean ± SD (n = 2). **b,** Cell viability data for ZH-2-077 compared to matching nitrogen-delete analogs (**25**-**27**) in gastric carcinoma cells and noncancerous HEK293T cells. Data shown are mean ± SD (n = 2). **c,** TMT non-enriched proteomics shows expression of ACAT2 and other detected proteins are not changed in soluble proteomes from cells treated with ZH-2-077 and ZH-5-049-1 at 10 µM (*in situ,* 2 h). ACAT2 is highlighted in blue and was detected exclusively in the soluble fraction. The proteins represent the combined results from n = 3 independent biological replicates. **d,** Competitive SuPUR TMT-ABPP evaluation of target engagement and proteome-wide selectivity of ZH-2-077 and matching nitrogen-delete ZH-5-049-1 control in 22Rv1 cells (*in situ,* 2 h). ZH-2-077 treatment results in significant liganding (CR<0.5. p<0.01) of ACAT2 Y237 in a site-specific and proteome-wide selective manner in soluble proteome from treated 22Rv1 cells. The AHL-PuP-2 modified peptides represent the combined results from n = 2 independent biological replicates. **e,** TMT non-enriched proteomics shows expression of ACAT2 and other proteins are not changed in 22Rv1 cells treated with ZH-2-077 or ZH-5-049-1 at 10 µM (*in situ*, 2 h). The proteins represent the combined results from n = 3 independent biological replicates. **f,** The expression of ACAT2 gene in tissues from different cancers (red) compared to normal tissues (gray). ACAT2 was significantly upregulated in cervical squamous cell carcinoma, endocervical adenocarcinoma (CESC) and prostate adenocarcinoma (PRAD) compared with normal tissues. Analysis of the ACAT2 gene reveals its association with various types of cancer and its impact on patient survival. Elevated ACAT2 expression is associated with a poorer overall survival (OS) rate. The results were generated using GEPIA 2.0 online tools.^63^ **g,** Mutation of ACAT2 Y237 to phenylalanine (Y237F) results in near complete loss of AHL-PuP-2 (25 µM) labeling as measured by gel-based ABPP and supports the hyper-reactive nature of this site (n = 3). **h,** Gel-based competitive ABPP analysis of ZH-2-077 and matching nitrogen-delete analogs ZH-5-049-1 and ZH-5-051 (*in vitro*, 1 h) shows the importance of purine recognition for ZH-2-077 activity. Data shown are mean ± SD (n = 3). **i,** The reactivities of SuPUR probes with ACAT2 were determined by gel-based ABPP in recombinant ACAT2-HEK293T soluble proteomes (*in vitro*, 1 h). SuPUR probes exhibited regioselective binding toward ACAT2, with *N7-*substituted AHL-PuP-2 showing more prominent labeling activity compared with matching *N9*-counterpart (ZH-2-087). p values were analyzed by t-test, **p<0.01. Data shown are mean ± SD (n = 3). **j,** ACAT2 catalyzes the reversible condensation of two acetyl-CoA molecules to form acetoacetyl-CoA and free CoA. The fluorescence-based assay was established using CoA Green as a probe to quantify the amount of released CoA; CoA Green reacts specifically with the thiol (–SH) group on CoA, generating a fluorescent signal that serves as a readout for ACAT2 enzymatic activity. To assess the robustness of the CoA-based fluorescence assay, the Z’ factor was determined using multiple biological replicates. A Z’ factor of 0.6 indicated good assay quality, making it suitable to screen for ACAT2 inhibitors.

**Extended data Fig. 8.**
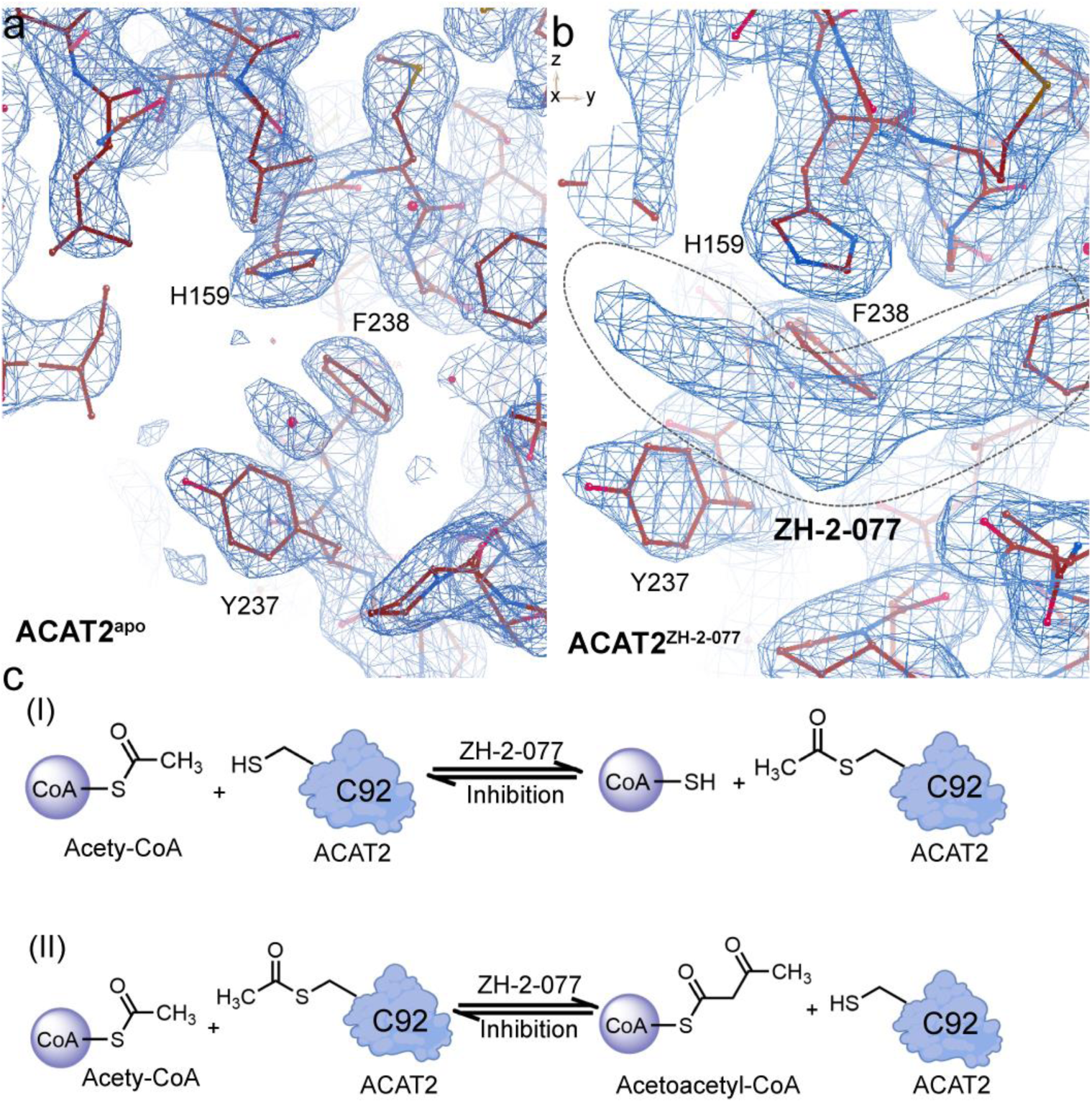
Co-crystal ACAT2^ZH-2-0^^77^ complex and ACAT2^apo^ protein analysis. **a** and **b,** The crystal structures of (**a**) ACAT2^apo^ and (**b**) ACAT2^ZH-2-077^. Comparison of the electron density maps near the active site between ACAT2^apo^ and ACAT2^ZH-2-077^ reveals an additional electron density in ACAT2^ZH-2-0^^77^, suggesting the presence of a bound ligand ZH-2-077. The composite (2*mF_o_* – *DF_c_*) omit map colored blue was contoured at 1.0 σ. This figure was generated using COOT. **c,** The acetyltransferase reaction mechanism involves the transfer of the acetyl group from acetyl-CoA to C92, followed by a Claisen condensation step.^65^ More details are provided in Supplementary **Table 5**.

## References

1. Burnstock, G. Purinergic signalling and disorders of the central nervous system. Nat. Rev. Drug Discov. 7, 575–590 (2008).

2. Migaud, M.E., Ziegler, M. & Baur, J.A. Regulation of and challenges in targeting NAD(+) metabolism. Nat. Rev. Mol. Cell Biol. 25, 822–840 (2024).

3. Longarini, E.J. & Dauben, H. Deciphering ADP-ribosylation signalling. Nat. Rev. Mol. Cell Biol. 25, 3 (2024).

4. Guertin, D.A. & Wellen, K.E. Acetyl-CoA metabolism in cancer. Nat. Rev. Cancer 23, 156–172 (2023).

5. An, S., Kumar, R., Sheets, E.D. & Benkovic, S.J. Reversible compartmentalization of de novo purine biosynthetic complexes in living cells. Science 320, 103–106 (2008).

6. Pareek, V., Tian, H., Winograd, N. & Benkovic, S.J. Metabolomics and mass spectrometry imaging reveal channeled de novo purine synthesis in cells. Science 368, 283–290 (2020).

7. Knapp, M., Bellamacina, C., Murray, J.M. & Bussiere, D.E. Targeting cancer: the challenges and successes of structure-based drug design against the human purinome. Curr. Top Med. Chem. 6, 1129–1159 (2006).

8. Murray, J.M. & Bussiere, D.E. Targeting the purinome. Methods Mol. Biol. 575, 47–92 (2009).

9. Haystead, T.A. The purinome, a complex mix of drug and toxicity targets. Curr. Top Med. Chem. 6, 1117–1127 (2006).

10. Patricelli, M.P. et al. Functional interrogation of the kinome using nucleotide acyl phosphates. Biochemistry 46, 350–358 (2007).

11. Ware, T.B. et al. Reprogramming fatty acyl specificity of lipid kinases via C1 domain engineering. Nat Chem Biol 16, 170–178 (2020).

12. Huang, M. & Wang, Y. Global and Targeted Profiling of Gtp-Binding Proteins in Biological Samples by Mass Spectrometry. Mass Spectrom Rev. 215–235 (2021).

13. Yu, M., de Carvalho, L.P., Sun, G. & Blanchard, J.S. Activity-based substrate profiling for Gcn5-related N-acetyltransferases: the use of chloroacetyl-coenzyme A to identify protein substrates. J. Am. Chem. Soc. 128, 15356–15357 (2006).

14. Hwang, Y., et al. A selective chemical probe for coenzyme A-requiring enzymes. Angew. Chem. Int. Ed. Engl. 46, 7621-7624 (2007).

15. Montgomery, D.C., Sorum, A.W. & Meier, J.L. Chemoproteomic profiling of lysine acetyltransferases highlights an expanded landscape of catalytic acetylation. J. Am. Chem. Soc. 136, 8669–8676 (2014).

16. Montgomery, D.C., Sorum, A.W., Guasch, L., Nicklaus, M.C. & Meier, J.L. Metabolic Regulation of Histone Acetyltransferases by Endogenous Acyl-CoA Cofactors. Chem. Biol. 22, 1030–1039 (2015).

17. Horning, B.D. et al. Chemical Proteomic Profiling of Human Methyltransferases. J. Am. Chem. Soc. 138, 13335–13343 (2016).

18. Chen, D. et al. Chemoproteomic Study Uncovers HemK2/KMT9 As a New Target for NTMT1 Bisubstrate Inhibitors. ACS Chem. Biol. 16, 1234–1242 (2021).

19. Janku, F., Yap, T.A. & Meric-Bernstam, F. Targeting the PI3K pathway in cancer: are we making headway? Nat. Rev. Clin. Oncol. 15, 273–291 (2018).

20. Denz, C.R. et al. Abstract ND06: First disclosure of AZD8421, a highly selective CDK2 inhibitor to address resistance to CDK4/6 inhibitors in breast and CCNE1-high cancers. Cancer Res. 84, ND06–ND06 (2024).

21. Denessiouk, K.A., Rantanen, V.V. & Johnson, M.S. Adenine recognition: a motif present in ATP-, CoA-, NAD-, NADP-, and FAD-dependent proteins. Proteins 44, 282–291 (2001).

22. Carugo, O. & Argos, P. NADP-dependent enzymes. II: Evolution of the mono- and dinucleotide binding domains. Proteins 28, 29–40 (1997).

23. Crespo-Hernández, C.E. et al. Electronic and structural elements that regulate the excited-state dynamics in purine nucleobase derivatives. J. Am. Chem. Soc. 137, 4368–4381 (2015).

24. Kimsey, I. & Al-Hashimi, H.M. Increasing occurrences and functional roles for high energy purine-pyrimidine base-pairs in nucleic acids. Curr. Opin. Struct. Biol. 24, 72–80 (2014).

25. Roundtree, I.A., Evans, M.E., Pan, T. & He, C. Dynamic RNA Modifications in Gene Expression Regulation. Cell 169, 1187–1200 (2017).

26. Xu, J. et al. Selective prebiotic formation of RNA pyrimidine and DNA purine nucleosides. Nature 582, 60–66 (2020).

27. Heindel, A.J. et al. Chemoproteomic capture of RNA binding activity in living cells. Nat. Commun. 14, 6282 (2023).

28. Ciancone, A.M. et al. Global Discovery of Covalent Modulators of Ribonucleoprotein Granules. J. Am. Chem. Soc. 145, 11056–11066 (2023).

29. Kim, G., Grams, R.J. & Hsu, K.L. Advancing Covalent Ligand and Drug Discovery beyond Cysteine. Chem. Rev. (2025).

30. Jones, L.H. Advances in sulfonyl exchange chemical biology: expanding druggable target space. Chem. Sci. (2025).

31. Hahm, H.S. et al. Global targeting of functional tyrosines using sulfur-triazole exchange chemistry. Nat. Chem. Biol. 16, 150–159 (2020).

32. Justin Grams, R., et al. Imidazoles are Tunable Nucleofuges for Developing Tyrosine-Reactive Electrophiles. Chembiochem 25, e202400382 (2024).

33. Rostovtsev, V.V., Green, L.G., Fokin, V.V. & Sharpless, K.B. A Stepwise Huisgen Cycloaddition Process: Copper(I)-Catalyzed Regioselective “Ligation” of Azides and Terminal Alkynes. Angew. Chem. Int. Ed. Engl. 41, 2596-2599 (2002).

34. Brulet, J.W., Borne, A.L., Yuan, K., Libby, A.H. & Hsu, K.L. Liganding Functional Tyrosine Sites on Proteins Using Sulfur-Triazole Exchange Chemistry. J. Am. Chem. Soc. 142, 8270–8280 (2020).

35. Ashburner, M. et al. Gene ontology: tool for the unification of biology. The Gene Ontology Consortium. Nat. Genet. 25, 25–29 (2000).

36. Gene Ontology, C., et al. The Gene Ontology knowledgebase in 2023. Genetics 224 (2023).

37. Moore, A.R., Rosenberg, S.C., McCormick, F. & Malek, S. RAS-targeted therapies: is the undruggable drugged? Nat. Rev. Drug Discov. 19, 533–552 (2020).

38. Bugter, J.M., Fenderico, N. & Maurice, M.M. Mutations and mechanisms of WNT pathway tumour suppressors in cancer. Nat. Rev. Cancer 21, 5–21 (2021).

39. Kanehisa, M. & Goto, S. KEGG: kyoto encyclopedia of genes and genomes. Nucleic Acids Res. 28, 27–30 (2000).

40. Huang, T. et al. Chemoproteomic profiling of kinases in live cells using electrophilic sulfonyl triazole probes. Chem. Sci. 12, 3295–3307 (2021).

41. Knox, C. et al. DrugBank 6.0: the DrugBank Knowledgebase for 2024. Nucleic Acids Res. 52, D1265–D1275 (2024).

42. Zdrazil, B. et al. The ChEMBL Database in 2023: a drug discovery platform spanning multiple bioactivity data types and time periods. Nucleic Acids Res. 52, D1180–d1192 (2024).

43. Kissinger, C.R. et al. Crystal structure of human ABAD/HSD10 with a bound inhibitor: implications for design of Alzheimer’s disease therapeutics. J. Mol. Biol. 342, 943–952 (2004).

44. Wang, X., Zhou, J. & Xu, B. Engaging an engineered PARP-2 catalytic domain mutant to solve the complex structures harboring approved drugs for structure analyses. Bioorg. Chem. 160, 108471 (2025).

45. Ogden, T.E.H. et al. Dynamics of the HD regulatory subdomain of PARP-1; substrate access and allostery in PARP activation and inhibition. Nucleic Acids Res. 49, 2266–2288 (2021).

46. Wu, H., Min, J., Zeng, H., Loppnau, P., Weigelt, J., Sundstrom, M., Arrowsmith, C.H., Edwards, A.M., Bochkarev, A., Plotnikov, A.N. Crystal Structure of GNPNAT1. (2006).

47. Zuhl, A.M. et al. Competitive activity-based protein profiling identifies aza-β-lactams as a versatile chemotype for serine hydrolase inhibition. J. Am. Chem. Soc. 134, 5068–5071 (2012).

48. Lajkiewicz, N.J., Cognetta, A.B., 3rd, Niphakis, M.J., Cravatt, B.F. & Porco, J.A., Jr. Remodeling natural products: chemistry and serine hydrolase activity of a rocaglate-derived β-lactone. J. Am. Chem. Soc. 136, 2659–2664 (2014).

49. Liu, Y., Patricelli, M.P. & Cravatt, B.F. Activity-based protein profiling: the serine hydrolases. Proc. Natl. Acad. Sci. U S A 96, 14694–14699 (1999).

50. Bachovchin, D.A. et al. Superfamily-wide portrait of serine hydrolase inhibition achieved by library-versus-library screening. Proc. Natl. Acad. Sci. U S A 107, 20941–20946 (2010).

51. Lustbader, J.W. et al. ABAD directly links Abeta to mitochondrial toxicity in Alzheimer’s disease. Science 304, 448–452 (2004).

52. Morsy, A. & Trippier, P.C. Amyloid-Binding Alcohol Dehydrogenase (ABAD) Inhibitors for the Treatment of Alzheimer’s Disease. J. Med. Chem. 62, 4252–4264 (2019).

53. Hroch, L. et al. Synthesis and evaluation of frentizole-based indolyl thiourea analogues as MAO/ABAD inhibitors for Alzheimer’s disease treatment. Bioorg. Med. Chem. 25, 1143–1152 (2017).

54. Benek, O. et al. Development of submicromolar 17beta-HSD10 inhibitors and their in vitro and in vivo evaluation. Eur. J. Med. Chem. 258, 115593 (2023).

55. Cao, Y. et al. ABHD10 is an S-depalmitoylase affecting redox homeostasis through peroxiredoxin-5. Nat. Chem. Biol. 15, 1232–1240 (2019).

56. Bachovchin, D.A. & Cravatt, B.F. The pharmacological landscape and therapeutic potential of serine hydrolases. Nat. Rev. Drug Discov. 11, 52–68 (2012).

57. Yan, S.D. & Stern, D.M. Mitochondrial dysfunction and Alzheimer’s disease: role of amyloid-beta peptide alcohol dehydrogenase (ABAD). Int. J. Exp. Pathol. 86, 161–171 (2005).

58. Powell, A.J. et al. Recognition of structurally diverse substrates by type II 3-hydroxyacyl-CoA dehydrogenase (HADH II)/amyloid-beta binding alcohol dehydrogenase (ABAD). J. Mol. Biol. 303, 311–327 (2000).

59. Bhatta, A., Dienemann, C., Cramer, P. & Hillen, H.S. Structural basis of RNA processing by human mitochondrial RNase P. Nat. Struct. Mol. Biol. 28, 713–723 (2021).

60. Mu, H. et al. OmicShare tools: A zero-code interactive online platform for biological data analysis and visualization. Imeta 3, e228 (2024).

61. Zhou, S. et al. Subclinical ketosis leads to lipid metabolism disorder by downregulating the expression of acetyl-coenzyme A acetyltransferase 2 in dairy cows. J. Dairy Sci. 106, 9892–9909 (2023).

62. Pietrocola, F., Galluzzi, L., Bravo-San Pedro, J.M., Madeo, F. & Kroemer, G. Acetyl coenzyme A: a central metabolite and second messenger. Cell Metab. 21, 805–821 (2015).

63. Li, C., Tang, Z., Zhang, W., Ye, Z. & Liu, F. GEPIA 2021: integrating multiple deconvolution-based analysis into GEPIA. Nucleic Acids Res. 49, W242–w246 (2021).

64. Zhang, M. et al. ACAT2 suppresses the ubiquitination of YAP1 to enhance the proliferation and metastasis ability of gastric cancer via the upregulation of SETD7. Cell Death Dis. 15, 297 (2024).

65. Kursula, P., Sikkilä, H., Fukao, T., Kondo, N. & Wierenga, R.K. High resolution crystal structures of human cytosolic thiolase (CT): a comparison of the active sites of human CT, bacterial thiolase, and bacterial KAS I. J. Mol. Biol. 347, 189–201 (2005).

66. Ferguson, F.M. & Gray, N.S. Kinase inhibitors: the road ahead. Nat. Rev. Drug Discov. 17, 353–377 (2018).

67. Wang, Y. et al. Expedited mapping of the ligandable proteome using fully functionalized enantiomeric probe pairs. Nat. Chem. 11, 1113–1123 (2019).

68. Njomen, E. et al. Multi-tiered chemical proteomic maps of tryptoline acrylamide-protein interactions in cancer cells. Nat. Chem. 16, 1592–1604 (2024).

69. Chen, Y. et al. Direct mapping of ligandable tyrosines and lysines in cells with chiral sulfonyl fluoride probes. Nat. Chem. 15, 1616–1625 (2023).

70. Cisar, J.S., et al. Identification of ABX-1431, a Selective Inhibitor of Monoacylglycerol Lipase and Clinical Candidate for Treatment of Neurological Disorders. J. Med. Chem. 61, 9062–9084 (2018).

71. Cheviet, T., Lefebvre-Tournier, I., Wein, S. & Peyrottes, S. Plasmodium Purine Metabolism and Its Inhibition by Nucleoside and Nucleotide Analogues. J. Med. Chem. 62, 8365–8391 (2019).

## References

72. Otwinowski, Z. & Minor, W. Processing of X-ray diffraction data collected in oscillation mode. Methods Enzymol. 276, 307–326 (1997).

73. Vagin, A. & Teplyakov, A. Molecular replacement with MOLREP. Acta Crystallogr. D. Biol. Crystallogr. 66, 22–25 (2010).

74. Emsley, P., Lohkamp, B., Scott, W.G. & Cowtan, K. Features and development of Coot. *Acta Crystallogr*. D. Biol. Crystallogr. 66, 486–501 (2010).

75. Murshudov, G.N., Vagin, A.A. & Dodson, E.J. Refinement of macromolecular structures by the maximum-likelihood method. Acta Crystallogr. D. Biol. Crystallogr 53, 240–255 (1997).

76. Li, Z. et al. In Situ Inhibitor Synthesis and Screening by Fluorescence Polarization: An Efficient Approach for Accelerating Drug Discovery. Angew. Chem. Int. Ed. Engl. 61, e202211510 (2022).

77. Cheng, Y. & Prusoff, W.H. Relationship between the inhibition constant (K1) and the concentration of inhibitor which causes 50 per cent inhibition (I50) of an enzymatic reaction. Biochem. Pharmacol. 22, 3099–3108 (1973).

78. Trott, O. & Olson, A.J. AutoDock Vina: improving the speed and accuracy of docking with a new scoring function, efficient optimization, and multithreading. J. Comput. Chem. 31, 455–461 (2010).

79. Eberhardt, J., Santos-Martins, D., Tillack, A.F. & Forli, S. AutoDock Vina 1.2.0: New Docking Methods, Expanded Force Field, and Python Bindings. J. Chem. Inf. Model. 61, 3891–3898 (2021).

80. Shin, M., Ware, T.B. & Hsu, K.L. DAGL-Beta Functions as a PUFA-Specific Triacylglycerol Lipase in Macrophages. Cell Chem. Biol. 27, 314–321.e315 (2020).

81. Gingrich, P.W., et al. canSAR 2024-an update to the public drug discovery knowledgebase. Nucleic Acids Res. 53, D1287–D1294 (2025).

82. Laskowski, R.A. SURFNET: a program for visualizing molecular surfaces, cavities, and intermolecular interactions. J. Mol. Graph. 13, 323–330, 307-328 (1995).

83. Lipinski, C.A., Lombardo, F., Dominy, B.W. & Feeney, P.J. Experimental and computational approaches to estimate solubility and permeability in drug discovery and development settings. Adv. Drug Deliv. Rev. 64, 4–17 (2012).

84. Veber, D.F. et al. Molecular properties that influence the oral bioavailability of drug candidates. J. Med. Chem. 45, 2615–2623 (2002).

85. Bekker, J. & Davis, J. Learning from positive and unlabeled data: a survey. Mach. Learn. 109, 719–760 (2020).

86. Xu, S. et al. Structural insight into the rearrangement of the switch I region in GTP-bound G12A K-Ras. Acta Crystallogr. D. Struct. Biol. 73, 970–984 (2017).

87. Kano, Y. et al. Tyrosyl phosphorylation of KRAS stalls GTPase cycle via alteration of switch I and II conformation. Nat. Commun. 10, 224 (2019).

88. Kuo, P.H., Chiang, C.H., Wang, Y.T., Doudeva, L.G. & Yuan, H.S. The crystal structure of TDP-43 RRM1-DNA complex reveals the specific recognition for UG- and TG-rich nucleic acids. Nucleic Acids Res. 42, 4712–4722 (2014).

89. Zong, Z., et al. Alanyl-tRNA synthetase, AARS1, is a lactate sensor and lactyltransferase that lactylates p53 and contributes to tumorigenesis. Cell 187, 2375–2392.e2333 (2024).

90. Graham, T.A., Ferkey, D.M., Mao, F., Kimelman, D. & Xu, W. Tcf4 can specifically recognize beta-catenin using alternative conformations. Nat. Struct. Biol. 8, 1048–1052 (2001).

91. van Veelen, W. et al. β-catenin tyrosine 654 phosphorylation increases Wnt signalling and intestinal tumorigenesis. Gut 60, 1204–1212 (2011).

92. Panne, D., Maniatis, T. & Harrison, S.C. An atomic model of the interferon-beta enhanceosome. Cell 129, 1111–1123 (2007).

